# SETD2 maintains nuclear lamina stability to safeguard the genome

**DOI:** 10.1101/2023.09.28.560032

**Authors:** Abid Khan, James M. Metts, Lucas C. Collins, C. Allie Mills, Kelin Li, Amanda L. Brademeyer, Brittany M. Bowman, M. Ben Major, Jeffrey Aubé, Laura E. Herring, Ian J. Davis, Brian D. Strahl

## Abstract

Histone methyltransferases play essential roles in the organization and function of chromatin. They are also frequently mutated in human diseases including cancer^1^. One such often mutated methyltransferase, SETD2, associates co-transcriptionally with RNA polymerase II and catalyzes histone H3 lysine 36 trimethylation (H3K36me3) – a modification that contributes to gene transcription, splicing, and DNA repair^2^. While studies on SETD2 have largely focused on the consequences of its catalytic activity, the non-catalytic functions of SETD2 are largely unknown. Here we report a catalysis-independent function of SETD2 in maintaining nuclear lamina stability and genome integrity. We found that SETD2, via its intrinsically disordered N-terminus, associates with nuclear lamina proteins including lamin A/C, lamin B1, and emerin. Depletion of SETD2, or deletion of its N-terminus, resulted in widespread nuclear morphology abnormalities and genome stability defects that were reminiscent of a defective nuclear lamina. Mechanistically, the N-terminus of SETD2 facilitates the association of the mitotic kinase CDK1 with lamins, thereby promoting lamin phosphorylation and depolymerization required for nuclear envelope disassembly during mitosis. Taken together, our findings reveal an unanticipated link between the N-terminus of SETD2 and nuclear lamina organization that may underlie how SETD2 acts as a tumor suppressor.

## Main

SETD2 is an evolutionarily conserved, nucleosomal-specific histone methyltransferase that associates with elongating RNA Polymerase II (RNAPII) and that catalyzes trimethylation of histone H3 lysine 36 (H3K36me3) in transcribed regions of the genome^3^. Studies have been focused on the function of SETD2’s interaction with RNAPII and how its methylation at H3K36 contributes to multiple chromatin-regulated processes including gene expression and DNA repair^4^. Like other sites of histone methylation, H3K36me3 functions by recruiting ‘reader’ proteins and/or complexes that possess specialized domains capable of recognizing H3K36 methylation^2, 4^. SETD2 is frequently mutated in human diseases including cancer^5^. Perhaps most recognized are the extensive mutations of SETD2 found in clear cell renal cell carcinomas (ccRCCs), which implicate SETD2 as a tumor suppressor^6^. Clear cell renal cell carcinomas are known for having an early structural chromosome abnormality wherein the 3p chromosomal arm is lost, thereby deleting one copy of *SETD2* and resulting in haploinsufficiency, a hallmark of ccRCC^7, 8^. Haploinsufficient *SETD2* tumors retain H3K36 methylation, yet they have drastic genome stability defects^9^. This finding has suggested that SETD2 performs important functions distinct from H3K36 methylation.

### SETD2 interacts with nuclear lamina proteins

All eukaryotic SETD2 share a common set of functional domains that includes an evolutionarily conserved SET domain, which mediates H3K36me3, and a C-terminal SRI domain that interacts with RNAPII (Extended Data Fig. 1a). Unlike SETD2 in less complex organisms such as yeast, mammalian SETD2 enzymes possess an extended N-terminal domain that is poorly characterized and enriched with intrinsically disordered regions (IDRs), most notably within the first 500 amino acids (Extended Data Fig. 1a). Because genetic studies have shown that the N-terminus of SETD2 is dispensable for H3K36me3 production in cells^10^, most studies have focused on the C-terminal half of SETD2 that mediates H3K36 methylation. Yet, studies by Bhattacharya et al. indicate that the N-terminus of SETD2 contributes to the regulation of SETD2’s stability, implying that this region is endowed with important regulatory functions^11, 12^. Additionally, recent studies have defined new protein-protein interaction motifs in the C-terminal half of SETD2^13^. Collectively, these findings suggest that other important regulatory mechanisms of SETD2 remain to be discovered.

To further understand how SETD2 contributes to chromatin biology, we extensively mapped the SETD2 protein-protein interactome. We employed a proximity-based biotinylation method using the ascorbate peroxidase, APEX2, which captures both stable and transient interactions^14^. A doxycycline inducible, full-length SETD2-APEX2 fusion construct was stably integrated into immortalized normal human kidney epithelial cells (HKC) and proximity labeling was performed, followed by mass spectrometry (Fig. 1a). Initial analysis yielded 1876 proteins that were significantly enriched over control (*P* < 0.05, Supplementary Table 1). Among these proteins were known interactors of SETD2 such as RNAPII subunits (e.g., POLR2A, POLR2B), thereby validating the robustness of the technique. Gene ontology classification of the SETD2 interactors (log2FC > 2, *P* < 0.05) revealed additional expected proteins involved in mRNA processing, transcription, and DNA repair (Fig. 1c, Supplementary Table 2). Unexpectedly, these analyses also showed a significant enrichment of proteins involved in nuclear envelope functions (Fig. 1c). Surveying the list of proteins in this group, we observed numerous nuclear lamina-associated proteins including lamin A/C, lamin B1, lamin B2, and emerin (Fig. 1b). SETD2-lamina association seemed paradoxical because lamins are located at the nuclear periphery where they typically tether heterochromatin to the lamina^15^, whereas SETD2 is known to associate with actively transcribed genes which are generally located towards the nuclear interior^16^. Nonetheless, we validated the SETD2-lamina interactions by performing immunoprecipitation of either endogenous SETD2 (Fig. 1d) or an exogenous, Halo-Flag tagged SETD2, followed by immunoblotting for endogenous lamina-associated proteins (Fig. 1e). Reverse co-immunoprecipitation using lamin A/C antibodies confirmed the interaction with SETD2 (Fig. 1f).

**Fig. 1.**
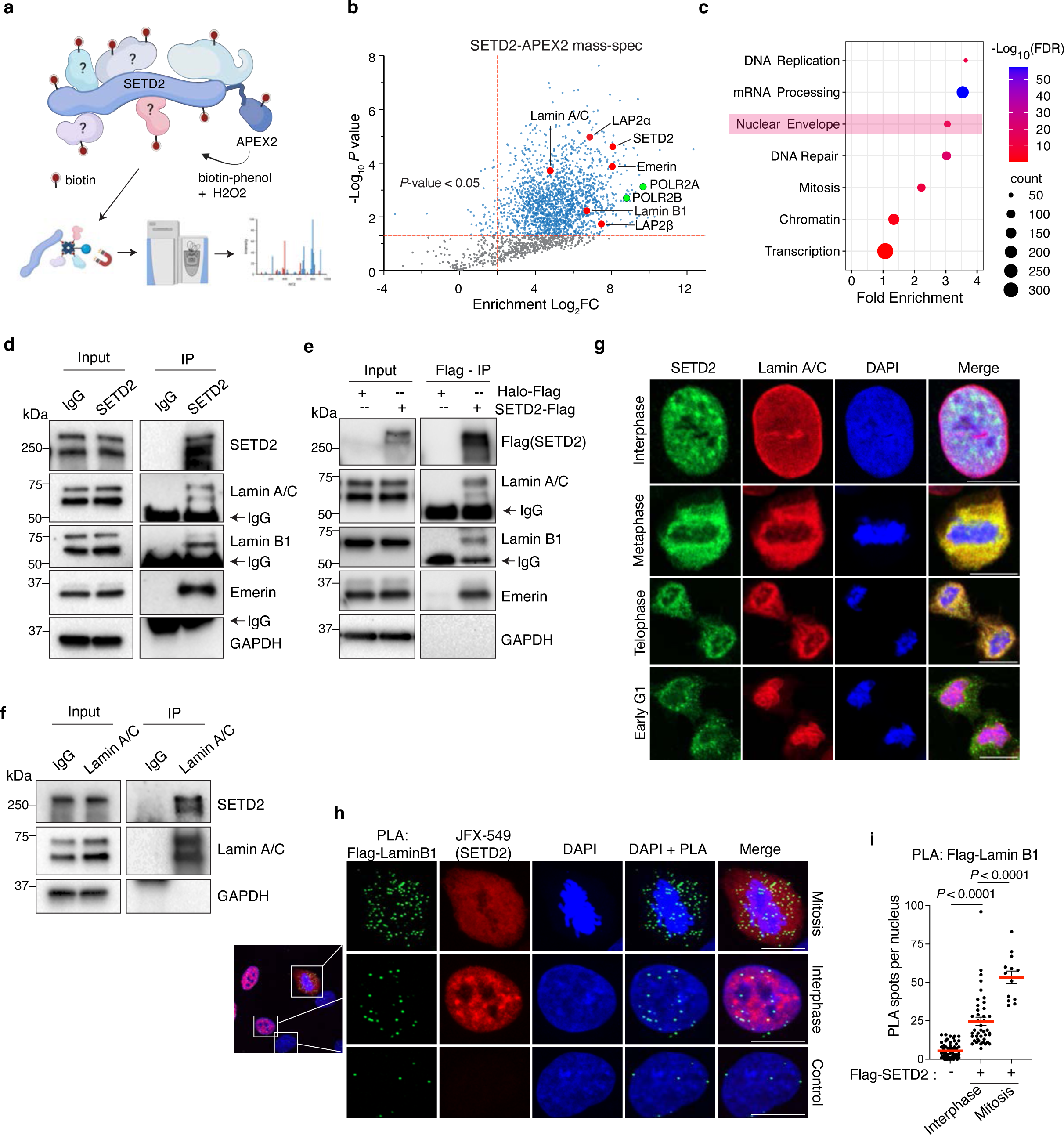
SETD2 interacts with nuclear lamina proteins. **a**, Schematic of SETD2-APEX2 and the experimental approach of the proximity biotinylation method. **b,** Volcano plot of SETD2-APEX2 mass spectrometry analysis, with nuclear lamina-associated proteins highlighted in red and RNAPII subunits in green. The x axis shows enrichment (Log2FC) of proteins in SETD2-APEX2-expressing (DoxON) cells compared with control cells (DoxOFF); the y-axis shows significance of enrichment. **c,** DAVID Gene Ontology (GO) analysis of mass-spec hits from 1b, defined as proteins with Log2FC>1 and p-value<0.05. Nuclear envelope is highlighted in pink. X-axis is fold enrichment and y-axis is significance (-logFDR). **d,** Immunoblot analyses for lamin proteins following immunoprecipitation of endogenous SETD2 with IgG as a control. **e,** Immunoblot analyses for endogenous lamin proteins following immunoprecipitation of exogenous Flag-Halo tagged SETD2 using anti-Flag antibodies with Halo-Flag as control. **f,** Immunoblot analyses for endogenous SETD2 upon lamin A/C immunoprecipitation with IgG as a control. **g,** Co-immunofluorescence of cells expressing exogenous SETD2 immunostained with anti-SETD2 (green) and anti-Lamin A/C (red) antibodies at the indicated cell cycle phase. **h,** Proximity ligation assay (PLA) with cells expressing exogenous Flag-Halo tagged SETD2 using anti-Flag and anti-Lamin B1 antibodies. PLA pairs are in green; SETD2 (red) visualized using fluorescent Halo ligand JFX-549. Control cells lack Flag-Halo-SETD2 and were a mixture of interphase and mitotic cells. **i,** Quantitative analysis of PLAs between Flag and Lamin B1 shown in h. Data are mean ± s.e.m. DNA was counterstained with DAPI. All scale bars are 10 µm.

The association of SETD2 with lamina proteins impelled us to determine when and where the interactions occur. As expected, co-immunofluorescence experiments revealed little overlap between SETD2 and lamins during interphase (Fig. 1g). In stark contrast, however, we observed strong co-localization of SETD2 with lamins in cells undergoing mitosis, which suggested that SETD2 functions with lamina proteins during this phase of the cell cycle when the nuclear envelope disassembles and lamins depolymerize^17^ (Fig. 1g and Extended Data Fig. 1b). To further examine the potential for *in vivo* interaction between SETD2 and lamins, we performed proximity-ligation assays (PLA) using a Flag antibody to detect Halo-Flag tagged SETD2 and lamin B1 antibody, which showed a significant increase in interaction frequency in mitotic cells compared with interphase cells (Fig. 1h, i). PLA experiments with antibodies toward endogenous SETD2 and lamin A/C also showed a similar trend of SETD2 association with lamins during mitosis (Extended Data Fig. 1c, d). Notably, the difference between interphase and mitosis interactions was lower with SETD2-lamin A/C pairs compared with SETD2-lamin B1 pairs. This difference may be due to the fact that lamin A/C is reported to occupy active genes and that some of the interaction between SETD2-lamin A/C could have occurred at genic regions during interphase^18^. In sum, these findings revealed an unsuspected link between SETD2 and nuclear lamina-associated proteins during mitosis, a link that demanded further examination.

### SETD2 loss leads to nuclear lamina defects

The nuclear lamina is a filamentous meshwork of lamin proteins that lies underneath the nuclear envelope. Lamin proteins homo-polymerize in a head-to-tail orientation to form filaments^17^. These filaments make contacts with large regions of heterochromatin to form stable structures known as lamina associated domains (LADs)^15^. By acting as a cushion to absorb external force, the lamina protects the LADs from external mechanical stress in the tissue microenvironment, thereby maintaining genome stability^19–21^. Indeed, tissue stiffness is directly proportional to lamin expression, which indicates that lamins provide tensile strength to the nuclei^22, 23^. Consistently, mutations or variations in lamin expression result in increased nuclear dysmorphia and genome instability in human diseases, particularly progeria and several cancers^17^.

To investigate the functional significance of SETD2-lamin interaction, we depleted SETD2 using a doxycycline inducible shRNA in HKC cells; we then performed confocal microscopy of these cells stained for lamin(s). Immunofluorescence experiments with SETD2 knockdown (SETD2-KD) cells revealed widespread nuclear abnormalities with a spectrum of different phenotypes, ranging from invaginations of lamina to micronuclei and blebbing, all hallmarks of a defective nuclear lamina (Fig. 2a). Quantification of the individual phenotypes showed that 20.53% of SETD2-KD cells had one or more micronuclei compared with 6.2% of control cells (Fig. 2b). Similarly, 20.3% SETD2-KD cells exhibited blebs as compared with 7.12% for control cells (Fig. 2d). In addition, we observed a subset of SETD2-KD cells (1.7%) that exhibited interphase bridges (Fig. 2f). To quantify invagination and nuclear shape, we adapted the CellProfiler pipeline designed to quantify such nuclear abnormalities^24^. Automated image analyses using CellProfiler showed a significant increase in the nuclear internal lamin B1 area, indicative of invagination of the lamina (Fig. 2c). Similarly, we observed a decrease in nuclei “roundness” (4πArea/perimeter^2^), which suggested an increase in misshaped nuclei in SETD2-KD cells (Fig. 2e). Notably, these phenotypes were not mutually exclusive because we observed a significant fraction of cells that exhibited more than one nuclear morphology defect (examples shown in Extended data 2a). In total, we found that nearly 40% of SETD2-KD cells had one or more nuclear defects (Fig. 2g).

**Fig. 2.**
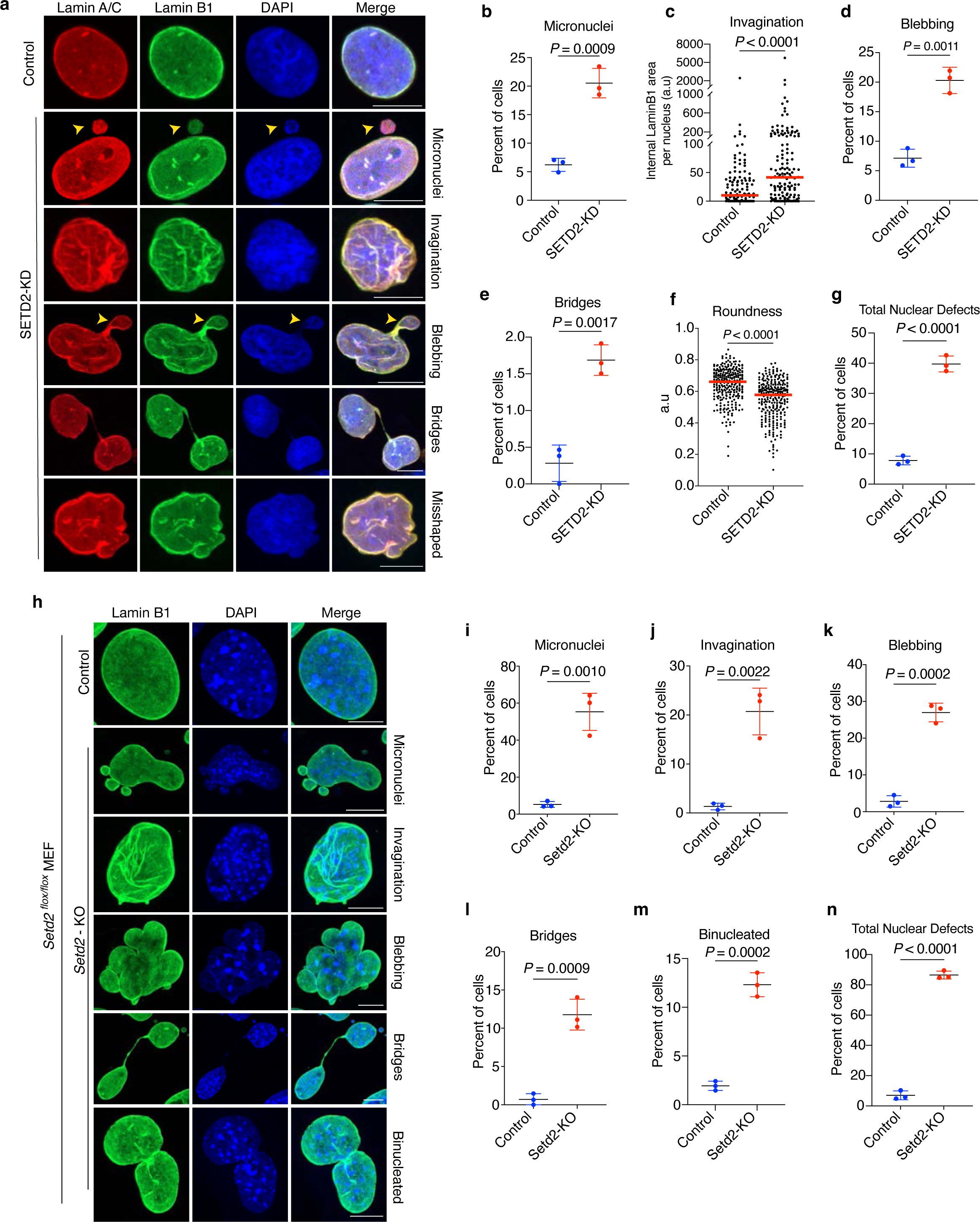
SETD2 loss leads to nuclear lamina defects. **a**, Representative immunofluorescence images of HKC cells immunostained with anti-Lamin A/C (red) and anti-Lamin B1 (green) antibodies in control and SETD2-depleted cells; indicated phenotypes are shown, and wider fields showing multiple cell examples are presented in Extended Fig. 2a. **b,** Quantification of cells having one or more micronuclei in control (DoxOFF) and SETD2-depleted (DoxON) cells. Two-tailed unpaired t-test of three independent biological replicates. Data are mean ± s.d. **c,** Quantification of invagination of lamina as internal Lamin B1 area per nucleus in control (n = 127) and SETD2-depleted cells (n = 146). Two-tailed non-parametric Mann-Whitney *U* test was performed; red bar indicates median. **d,** Quantification of cells exhibiting blebbing from three independent replicates. Two-tailed unpaired *t-*test was performed from three independent biological replicates; data shown are mean ± s.d. **e,** Quantification of nuclei Roundness (4πArea/perimeter^2^) in control (n = 272) and SETD2-depleted cells (n = 272). Two-tailed non-parametric Mann-Whitney *U* test was performed; red bar indicates median. **f,** Quantification of cells with interphase bridges from three independent replicates. Two-tailed unpaired *t-*test was performed from three independent biological replicates; data are mean ± s.d. **g,** Quantification of all nuclear defects from three independent replicates. Two-tailed unpaired *t-*test was performed from three independent biological replicates; data are mean ± s.d. **h,** Representative images of MEF cells immunostained with anti-lamin B1 (green) antibody in control (ethanol) and *Setd2* KO (tamoxifen) cells exhibiting indicated phenotypes. Individual cell examples are shown, and wider field views are presented Extended Data Fig. 2b. **i,** Quantification of cells having one or more micronuclei in control and *Setd2* KO cells. Two-tailed unpaired t-test on data from three independent biological replicates. Data shown are mean ± s.d. **j,** Quantification of cells exhibiting invagination of lamina in control and *Setd2* KO cells. Two-tailed unpaired t-test on data from three independent biological replicates. Data are mean ± s.d. **k,** Quantification of cells exhibiting blebbing in control and *Setd2* KO cells. Two-tailed unpaired t-test on three independent biological replicates. Data are mean ± s.d. **l,** Quantification as percent of cells having interphase bridges in control and *Setd2* KO cells. Two-tailed unpaired t-test on data from three independent biological replicates. Data are mean ± s.d. **m,** Quantification of binucleated control or *Setd2* KO cells. Two-tailed unpaired t-test on data from three independent biological replicates. Data are mean ± s.d. **n,** Quantification of all nuclear defects from control or *Setd2* KO cells. Two-tailed unpaired t-test on data from three independent biological replicates. Data are mean ± s.d. DNA was counterstained with DAPI. All scale bars are 10 µm.

The unexpected finding that SETD2 is required for nuclear lamina integrity prompted us to determine whether this requirement exists in other cell types. We used a mouse embryonic fibroblast (MEF) model system that harbored homozygous floxed alleles of *Setd2*, which enabled conditional knock-out upon 4-hydroxytamoxifen treatment^9^. Strikingly, tamoxifen treatment for two days led to profound nuclear defects, far more severe than the defects observed in *Setd2*-depleted HKC cells (Fig. 2h). The severity of the phenotypes and gross nuclear dysmorphia prevented us from using CellProfiler for automated image analysis. Manual tabulation of the phenotypes showed that 55.28% of *Setd2*-KO cells had one or more micronuclei compared with 5.25% of control cells (Fig. 2i). Additionally, nearly 21% of *Setd2*-KO cells exhibited invaginations and 27% had blebs as opposed to 1.4% and 2.8% of control cells, respectively (Fig. 2j, k). Interphase bridges were also observed in 12% *Setd2*-KO cells compared with 0.7% in control cells (Fig 2l), and 12.3% *Setd2*-KO cells were binucleated compared with 2% of control cells (Fig. 2m). In total, 86.47% of *Setd2*-KO cells exhibited one or more nuclear lamina phenotypes (Fig. 2n). Together, these data demonstrated a conserved function of SETD2 in the maintenance of nuclear lamina integrity and genome stability.

### Absence of SETD2 impairs lamin phosphorylation during G2/M

Because SETD2-lamin interaction occurs predominantly during mitosis, we wanted to gain a more detailed temporal picture of this interaction during G2-M. To this end, we performed live-cell confocal microscopy of control and SETD2-depleted cells that stably expressed emerald-lamin A. Prior to imaging, we synchronized cells using a double thymidine block and release method. Time-lapse imaging of control cells undergoing mitosis revealed that emerald-lamin A was completely depolymerized and showed homogenous signal by metaphase. In contrast, we observed a significant fraction (19.43%) of SETD2-depleted cells that exhibited partially depolymerized emerald-lamin A filaments that remained intact throughout mitosis (Fig. 3a, b and Extended data 3a). Additionally, we observed that lamina reassembly, post-mitosis, was slightly faster in SETD2-depleted cells compared with control cells (Fig. 3c). Thus, the absence of SETD2 changed the dynamics of nuclear envelope breakdown and reassembly during the G2-M phases of the cell cycle.

**Fig. 3.**
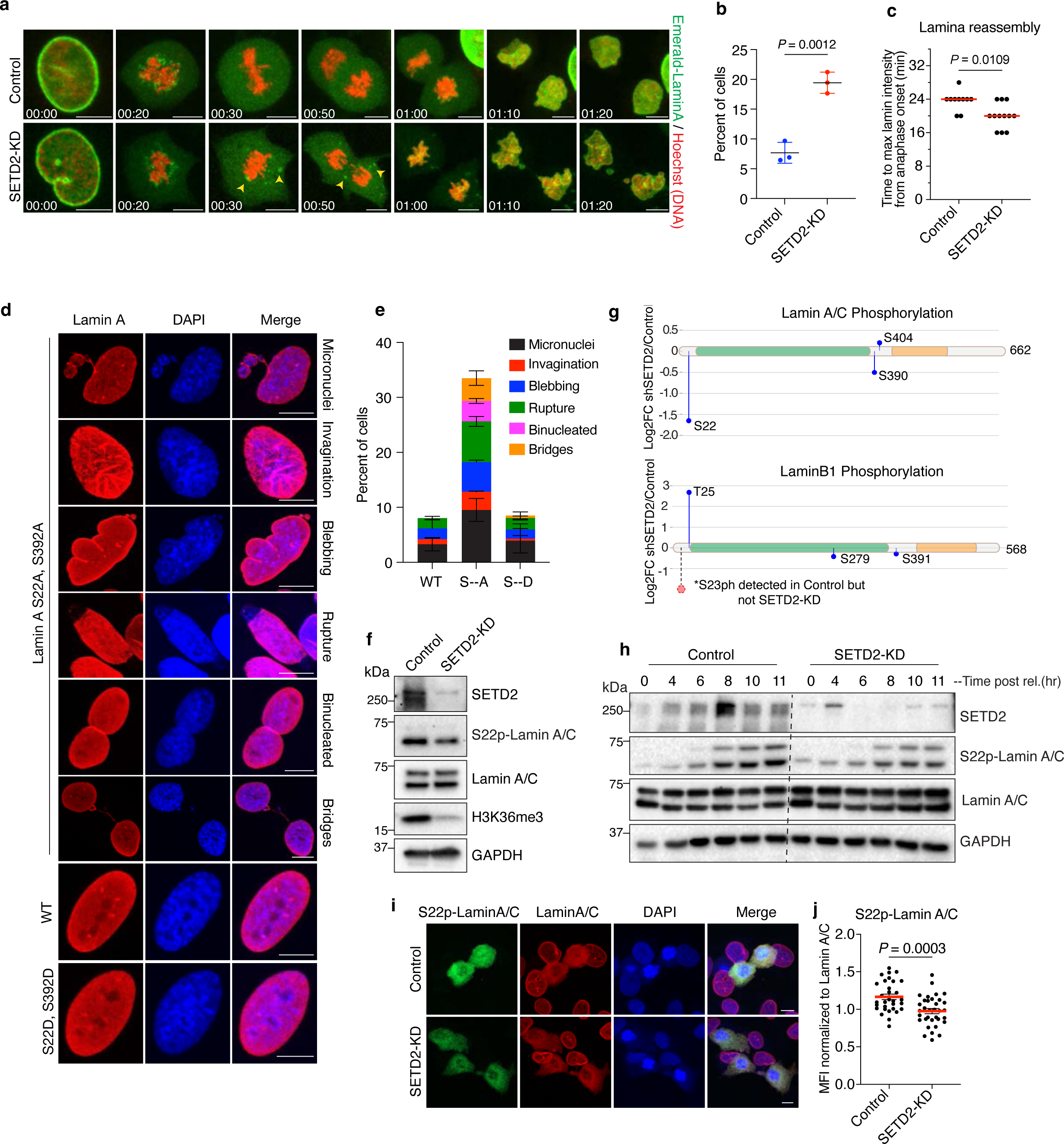
SETD2 loss impairs lamin phosphorylation during G2/M. **a**, Representative time-lapse images of control and SETD2-depleted HKC expressing Emerald-lamin A (green). DNA was labeled with SiR-Hoechst (red). Arrowheads point to partially depolymerized lamin filaments observed throughout mitosis. **b,** Quantification of cells having partially depolymerized Emerald-lamin A filaments from three independent replicates. Two-tailed parametric *t*-test was performed; data shown are mean ± s.d. **c,** Quantitative analysis of lamina reassembly from live-cell microscopy of Emerald-lamin A-expressing cells. Time taken to reach maximum intensity from anaphase onset was determined in control (n = 10) and SETD2-KD (n = 12) cells. Two-tailed non-parametric Mann-Whitney *U* test was performed; red bar indicates median. **d,** Representative images of indicated phenotypes from *LMNA* KO human fibroblasts cells expressing either WT, S->A (S22A; S392A) or S->D (S22D;S392D) mutant lamin A. Cells were immunostained with lamin A/C antibody (red) and DNA was counterstained with DAPI. **e,** Quantitative analysis of phenotypes shown in d. **f.** Immunoblot analysis of indicated proteins from control and SETD2-KD cells. **g,** Mass spectrometry analysis of immunoprecipitated lamin A/C and lamin B1 phosphorylation from three independent replicates. The x-axis shows position of amino acids of the indicated protein. Blue lollipop shows relative changes in phosphorylation level in SETD2-depleted cells compared to control. The y-axis is log2FC of shSETD2 over control. Dashed line and red circle indicate phosphorylation of serine 23 in lamin B1. Phospho-peptides of S23-lamin B1 were detected in all replicates of control but in neither of shSETD2 samples. **h,** Immunoblot analysis of control and shSETD2 cells synchronized by double thymidine block and released for indicated time points. **i,** Representative images of S22phos-lamin A/C (green) and pan lamin A/C (red) from control and SETD2 depleted cells. DNA was counterstained with DAPI. **j,** Quantitative analysis of immunofluorescence data from i. Mean fluorescence intensity (MFI) of S22ph-lamin A/C signal from mitotic cells normalized to MFI of pan lamin A/C signal in control (n = 31) and shSETD2 (n = 34) cells from two independent biological replicates. All scale bars are 10 µm.

Nuclear envelope disassembly is accomplished by the action of microtubule-dependent tearing of the nuclear envelope and subsequent depolymerization of filamentous lamin proteins^17^. Hyperphosphorylation of lamin proteins at several residues, particularly at several conserved serine residues (i.e., S22, S392 in lamin A/C and S23, S393 in lamin B1), drives depolymerization of lamin filaments and their detachment from chromatin^17, 25^. At the onset of anaphase, lamins are rapidly dephosphorylated by mitotic phosphatases such as PP1 and PP2A; dephosphorylation facilitates lamin polymerization back into filaments which then form the nuclear lamina and reestablish chromatin contacts in daughter nuclei^17, 25^. This cycle of phosphorylation-dependent depolymerization and re-polymerization is critical to the maintenance of genome integrity and proper nuclear morphology. In agreement with the requirement for this phosphorylation cycle, studies have shown that serine to alanine mutations of S22/S392 in lamin A or S23/S393 in lamin B1 lead to incomplete breakdown of lamin filaments, causing them to remain attached to chromosomes during mitosis^26, 27^.

The findings that SETD2 influenced the dynamics of lamin disassembly and reassembly prompted us to determine whether SETD2 might regulate lamina function by control of lamin phosphorylation. If this idea was correct, we would expect that mutations in lamin that prevent its phosphorylation should phenocopy the nuclear morphology defects observed upon depletion of SETD2. To test if this expectation was indeed the case, we employed *LMNA*-KO human fibroblasts that contained stably integrated doxycycline-inducible wild-type (WT) lamin A or lamin A that was mutated to S22A/S392A or S22D/S392D^28^. Analysis of these cells showed a striking similarity between defects observed in S22A/S392A lamin A mutant fibroblasts with the defects observed in SETD2-deficient HKC and MEFs (Fig. 3d, 3e and Extended data 3e). These similarities raised the possibility that SETD2 is directly involved in the dynamics of lamin phosphorylation during mitosis.

To determine whether SETD2 is important to the cycle of lamin phosphorylation, we initially examined the status of lamin A/C phosphorylation by immunoblot analysis in our WT and SETD2-KD HKC cells. Using an anti-S22phos-lamin A/C antibody to monitor lamin A/C phosphorylation, we observed that SETD2 depletion resulted in a decrease in overall S22 phosphorylation compared with the control (Fig. 3f). Because lamin phosphorylation occurs predominantly during the G2-M phases of the cell cycle, we wanted to ascertain whether the decrease in lamin A/C phosphorylation might be due simply to changes in the G2/M population of cells. Flow cytometry analyses did not reveal drastic changes in G2/M population that could explain the decrease in lamin A/C S22 phosphorylation in SETD2-depleted cells (Extended Fig. 3b, c). We note, and in agreement with our previous findings, these flow cytometry analyses did show that SETD2 depletion led to an increase in the G1 population of cells and an equal decrease in the S-phase population.

We next sought to quantitatively determine the effect of SETD2 depletion on lamin A/C phosphorylation and whether the phosphorylation of other lamins might be affected similarly to lamin A/C at S22. Accordingly, we immunoprecipitated lamin A/C and lamin B1 from control and SETD2-KD cells and performed quantitative mass spectrometry to measure the effects of SETD2 depletion on lamin phosphorylation. These analyses confirmed our immunoblot analyses by showing a significant decrease in S22 phosphorylation with SETD2 depletion compared to control, in addition to a partial decrease in S390 in lamin A/C. Importantly, phosphorylated peptides spanning S23 lamin B1 were detected in all control replicates but not in our SETD2-depleted samples, indicating a significant reduction in phosphorylation of S23 upon SETD2 depletion (Fig. 3g). Notably, not all phosphorylation sites in the lamins that we identified showed a decrease in SETD2 depleted cells; we also observed a significant increase in T25 phosphorylation in lamin B1, although the significance of this site of phosphorylation is unknown. Nonetheless, these data demonstrated that loss of SETD2 causes a decrease in lamin phosphorylation at specific sites that mediate lamin disassembly and reassembly.

The linkage of SETD2 to lamin phosphorylation during mitosis prompted us to examine the temporal relationship between lamin phosphorylation and cell cycle oscillation of SETD2 protein^29^. In earlier work, we found that SETD2 protein level peaked during the G2/M phase of the cell cycle, which suggested that SETD2 stability during G2/M might be functionally linked with the timing of lamin phosphorylation^25, 29^. To examine this possibility, we synchronized cells by double thymidine block and released into S phase, followed by immunoblot analyses of time points that spanned the G2 and M phase. Intriguingly, we found that the transient peak of SETD2 occurred at 8 hours post release, precisely coinciding with the onset of lamin phosphorylation in control cells, which occurs during late G2 phase of the cell cycle^25^. In SETD2-KD cells, and consistent with earlier results, lamin phosphorylation was decreased at all time points examined (Fig. 3h). Additionally, by quantitative analysis of immunofluorescence with an S22 phosho-specific lamin A/C antibody on nocodazole-arrested mitotic cells, we found a decrease in lamin A/C S22 phosphorylation in SETD2-depleted cells (Fig. 3i, j and Extended data 3e). Collectively, these data demonstrated that during G2-M phases SETD2 is required for proper lamin phosphorylation dynamics that governs nuclear envelope breakdown and lamina reassembly.

### A catalysis independent function of SETD2 in nuclear lamina integrity

Clear cell renal cell carcinomas often contain a heterozygous deletion of *SETD2* due to the loss of the 3p chromosome arm, yet they retain significant levels of H3K36 methylation^8, 9^. Cell-based models of *SETD2* haploinsufficiency also retain H3K36me3 in addition to harboring a variety of genome integrity defects that include micronuclei and chromosomal bridges^9^. Notably, several studies suggest SETD2 methylation of α-tubulin may be responsible for these genome stability defects^9, 30^. Given our findings that SETD2 associates with lamina proteins and that defects in lamin phosphorylation produce the same genome instability phenotypes as seen in our SETD2 KD or deletion cells, we wondered whether regulation of nuclear lamina integrity by SETD2 requires the methylation activity of SETD2. Thus, we treated HKC cells with a highly selective and potent inhibitor of SETD2 methyltransferase activity (EPZ-719; SETD2i)^31^. As expected, acute inhibition of SETD2 led to complete loss of H3K36me3 (Fig. 4a). Remarkably, inhibition of SETD2’s methylation activity did not lead to any significant nuclear defects (Fig. 4b, c). Furthermore, SETD2 inhibition did not reduce lamin phosphorylation levels (Fig. 4a). In complete contrast to the lack of inhibitor effect on nuclear integrity, acute depletion of SETD2 consistently resulted in both nuclear abnormalities in 38% of HKC cells and a decrease in lamin phosphorylation (Fig. 4a-c). To analyze these findings further, we used a MEF model system for *SETD2* haploinsufficiency, wherein one of the two *SETD2* genes was floxed, enabling conditional deletion of one copy of *SETD2.* As expected, deletion of one *SETD2* allele did not significantly alter H3K36me3 (Fig. 4d). Strikingly, however, loss of a single copy of *SETD2* was sufficient to drive nuclear lamina abnormalities and genome stability defects similar to those observed in *Setd2*-KO MEFs (Fig. 4e, f). Remarkably again, SETD2 inhibition by EPZ-719 did not result in any nuclear morphology or genome stability defects, even though H3K36me3 was almost completely ablated (Fig 4d-f). These data indicated two crucial facts. First, the methyltransferase activity of SETD2 is dispensable for the maintenance of the nuclear lamina and dispensable for prevention of a host of genome integrity defects previously thought to be associated with its catalytic function. Second, both alleles of *SETD2* are required to maintain nuclear lamina integrity. Thus, a precise intracellular concentration of SETD2 protein, and not its catalytic activity, is vital to the stability of the nucleus and integrity of the genome.

**Fig. 4.**
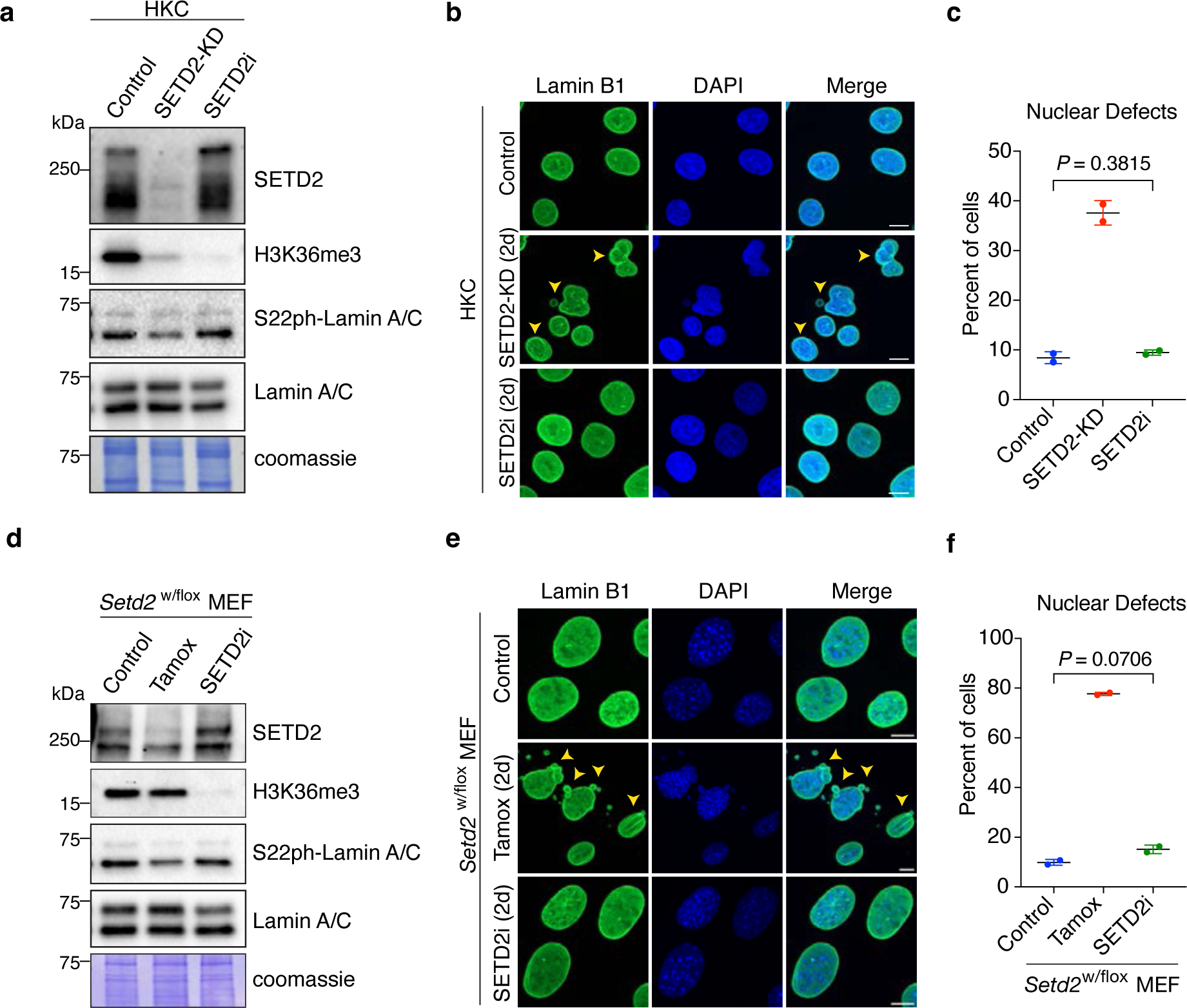
Nonenzymatic function of SETD2 in maintenance of nuclear integrity. **a**, Immunoblot analyses of indicated proteins from control, SETD2-KD and SETD2i (EPZ-719)-treated cells for 48 hr. **b,** Representative images of control, SETD2-KD and SETD2i-treated HKC cells immunostained with lamin B1 antibody. **c,** Quantitative analysis of nuclear defects from b. Two-tailed unpaired *t-*test was performed from two independent biological replicates; data are mean ± s.d. **d**, Immunoblot analyses of indicated proteins from control, Tamoxifen (*Setd2*-KO) and SETD2i (EPZ-719)-treated MEF cells for 48 hr. **e,** Representative images of control, *Setd2*-KO and SETD2i-treated HKC cells immunostained with lamin B1 antibody. **f** Quantitative analysis of nuclear defects from e. Two-tailed unpaired *t-*test was performed from two independent biological replicates; data shown are mean ± s.d. DNA was counterstained with DAPI. All scale bars are 10 µm.

### The N-terminus of SETD2 regulates nuclear lamina integrity

The foregoing experiments established that the catalytic activity of SETD2 was dispensable for maintenance of nuclear lamina integrity. This prompted us to ask which region or domain within SETD2 is contributing towards this function. To answer this question, we generated a doxycycline-inducible HKC cell system, wherein a stably integrated and doxycycline inducible shRNA expression construct would deplete endogenous SETD2 while simultaneously inducing the expression of either WT or various SETD2 mutant forms (Fig. 5a). We first performed immunoblot analyses, which confirmed previous reports that truncation of the N-terminus of SETD2 (tSETD2) can robustly restore H3K36me3. As expected, intact WT and the SRI domain mutant (SRI_mut_, R2510H), but not a SET domain mutant that was catalytically inactive (SET_mut_, R1625C), were also able to rescue H3K36 methylation. Immunoblot analysis of lamin A/C phosphorylation revealed that expression of either the WT, SET_mut_ or SRI_mut_ forms of SETD2 rescued lamin phosphorylation in SETD2-KD cells. In complete contrast to these findings, cells that expressed tSETD2 were deficient in lamin phosphorylation, similar to SETD2-KD cells. Thus, lamin phosphorylation was dependent on the N-terminus of SETD2 (Fig. 5b).

**Fig. 5.**
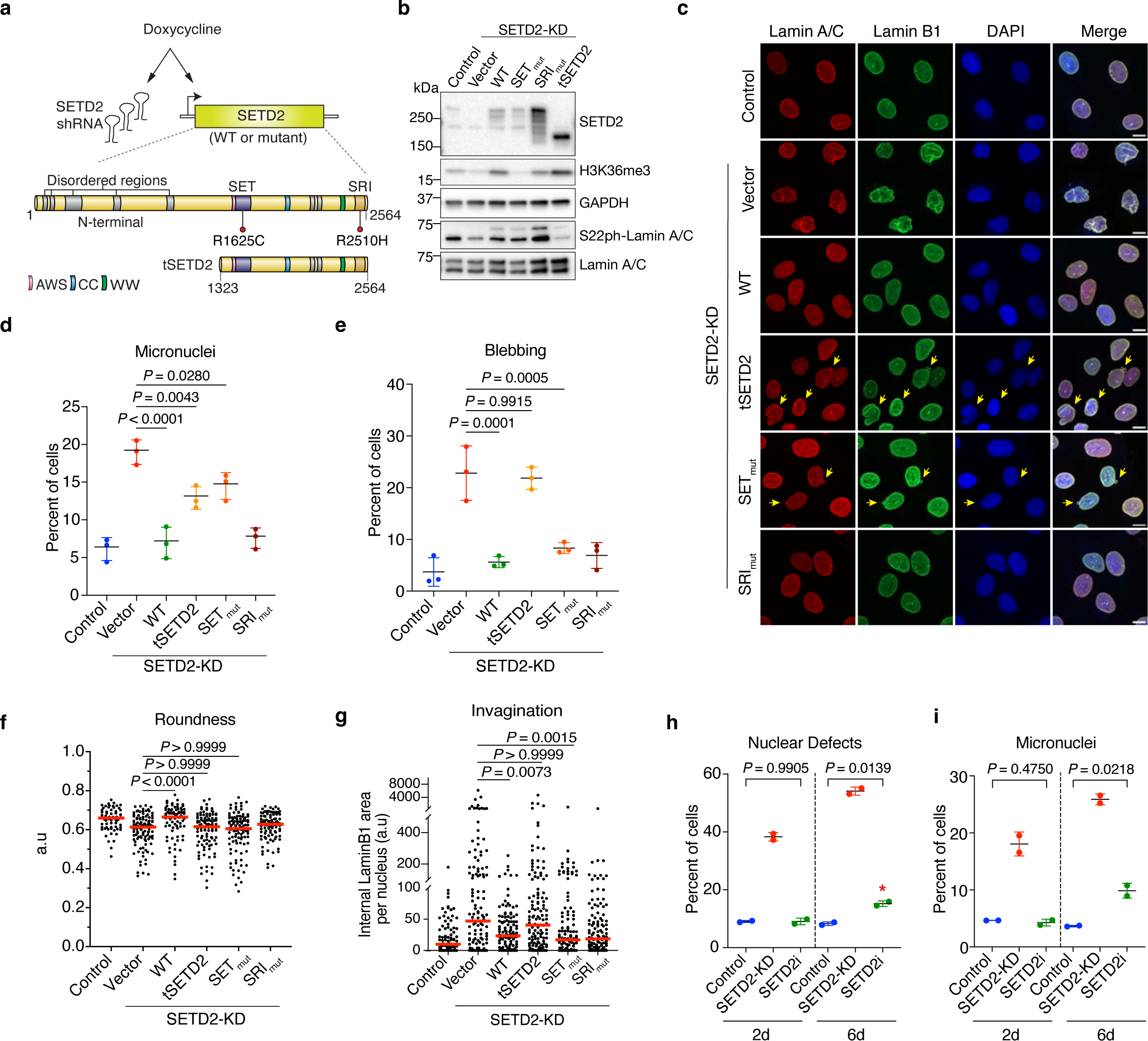
The N-terminus of SETD2 regulates nuclear lamina integrity. **a**, Schematic of doxycycline inducible shRNA and genetic complementation system in HKC cells. **b,** Immunoblot analyses of indicated proteins in control, SETD2-KD or SETD2-KD cells expressing either WT or mutant SETD2. **c**, Representative images of control or SETD2-KD cells expressing the indicated transgene, immunostained with lamin A/C and lamin B1 antibodies. DNA was counterstained with DAPI. Scale bar is 10µm. **d,** Quantification of cells containing one or more micronuclei. One-way ANOVA with Dunnett’s multiple comparisons test was performed on three independent biological replicates. Data shown are mean ± s.d. **e,** Quantification of cells with one or more blebs in the indicated samples. One-way ANOVA with Dunnett’s multiple comparisons test was performed on three independent biological replicates. Data shown are mean ± s.d. **f,** Quantification of nuclei Roundness (4πArea/perimeter^2^) in the indicated samples. Kruskal-Wallis with Dunn’s multiple comparisons test was performed on n > 100 cells from three biological replicates; red bar indicates median. **g,** Quantification of Invaginations in the indicated samples. Kruskal-Wallis with Dunn’s multiple comparisons test was performed on n > 100 cells from three biological replicates; red bar indicates median. **h**, Quantification of total nuclear defects in control, SETD2-KD or SETD2i-treated cells for 2d or 6d. Two-tailed unpaired *t-*test was performed from two independent biological replicates; data shown are mean ± s.d. **i**, Quantification of cells with micronuclei in control, SETD2-KD or SETD2i-treated cells for 2d or 6d. Two-tailed unpaired *t-*test was performed from two independent biological replicates; data shown are mean ± s.d.

To assess whether the nuclear defects observed in SETD2-KD cells were the same as the defects in tSETD2, we performed an immunofluorescence analysis of lamin proteins followed by confocal microscopy in WT and SETD2 mutants (Fig. 5c-g). Re-expression of WT SETD2 and the SRI_mut_ rescued all of the nuclear defects usually observed in SETD2-depleted cells. In contrast, expression of tSETD2 in SETD2-KD cells did not rescue the blebbing, invaginations, and roundness phenotypes. We note, however, that tSETD2 did lead to a partial rescue of micronuclei generation. Collectively, these data revealed that not only is the N-terminus of SETD2 required for lamin phosphorylation, but the N-terminus is essential to maintain nuclear lamina integrity and genome stability.

Our studies with the WT and mutant SETD2 forms also revealed a curious finding. Although the expression of the SET_mut_ in SETD2-KD cells rescued nuclear invaginations and blebbing, SET_mut_ did not completely rescue nuclear roundness and micronuclei generation (Fig. 5d, f). This finding seemed at odds with our acute SETD2 inhibition studies that showed SETD2 activity was dispensable to prevent nuclear morphology defects (Fig. 4a-c). However, a major difference between these experiments and the earlier assays was that the SETD2 complementation experiments required long time frames (6 days post dox treatment) to allow for full expression and rescue of H3K36me3 by our SETD2 constructs; previous knockdowns and our chemical inhibition studies were performed at 2 days, the earliest time at which significant SETD2 depletion and loss of H3K36me3 was observed. Therefore, we hypothesized that prolonged inhibition of SETD2 may lead to increased micronuclei generation. To examine this hypothesis, we performed a time course of SETD2 inhibition in our HKC cells for 2 or 6 days and compared these treatment times with a SETD2 knockdown. Consistently, SETD2-KD resulted in nuclear defects in approximately 40% of cells, whereas SETD2 inhibition with EPZ-719 did not result in nuclear defects at the 2-day time point (Fig. 5h and Extended data 4a). However, 6 days of SETD2 inhibition with EPZ-719 did lead to a small but significant increase in nuclear defects that were largely composed of micronuclei (Fig. 5h, i). Intriguingly, SETD2 depletion for 6 days also increased the overall percent of nuclear defects (from 37% [2d] to 53% [6d]), which indicated a compounding effect of prolonged SETD2 and H3K36me3 loss. We surmise that the phenotypes observed with prolonged SETD2 inhibition arose likely from secondary effects because a total loss of H3K36me3 at early time points did not have discernible nuclear or genome stability defects.

### SETD2 N-terminus interacts with lamins

Because tSETD2 did not rescue the nuclear and lamin phosphorylation abnormalities associated with depletion of SETD2, we sought to determine exactly how the N-terminus of SETD2 contributed to nuclear lamina stability. We employed APEX2-mediated proximity labeling to determine the protein-protein interaction differences between WT and tSETD2. We generated doxycycline-inducible APEX2 fusion constructs of WT and tSETD2 and stably integrated these constructs into our HKC cells for proximity labeling followed by mass spectrometry analysis. Initial analyses showed 1068 proteins significantly enriched in WT SETD2-expressing cells over tSETD2 expressing cells and 458 proteins significantly enriched in cells that expressed tSETD2 over WT (log2FC > 1, p<0.05) (Fig. 6a; Supplementary Table 1). Further analyses revealed that interaction of several nuclear lamina-associated proteins (i.e., lamin A/C, lamin B1 and emerin) were enriched with WT SETD2 compared with tSETD2 (Fig. 6a). To validate these results, we performed immunoprecipitation from cells that expressed either a Halo-Flag tagged WT SETD2 or tSETD2, followed by immunoblotting for endogenous lamin proteins. This analysis confirmed the mass spectrometry results and showed a striking difference between the interaction profile of WT SETD2 and tSETD2 with endogenous lamina-associated proteins (Fig. 6b). Specifically, WT SETD2 efficiently co-precipitated lamin A/C, lamin B1 and emerin, whereas tSETD2 did not associate with either lamin B1 or emerin. Intriguingly, we also observed differences between WT SETD2 and tSETD2 regarding lamin A and lamin C interaction; tSETD2 selectively associated with lamin C, whereas WT SETD2 associated with both lamin A and C. Because lamin C is localized at the nuclear interior during the early G1 phase of the cell cycle prior to its assembly into the nuclear envelope^32^, we suspect the differential preferences of tSETD2 and SETD2 for different lamins may underlie multiple SETD2-lamin functions (e.g., transcription vs nuclear lamina formation).

**Fig. 6.**
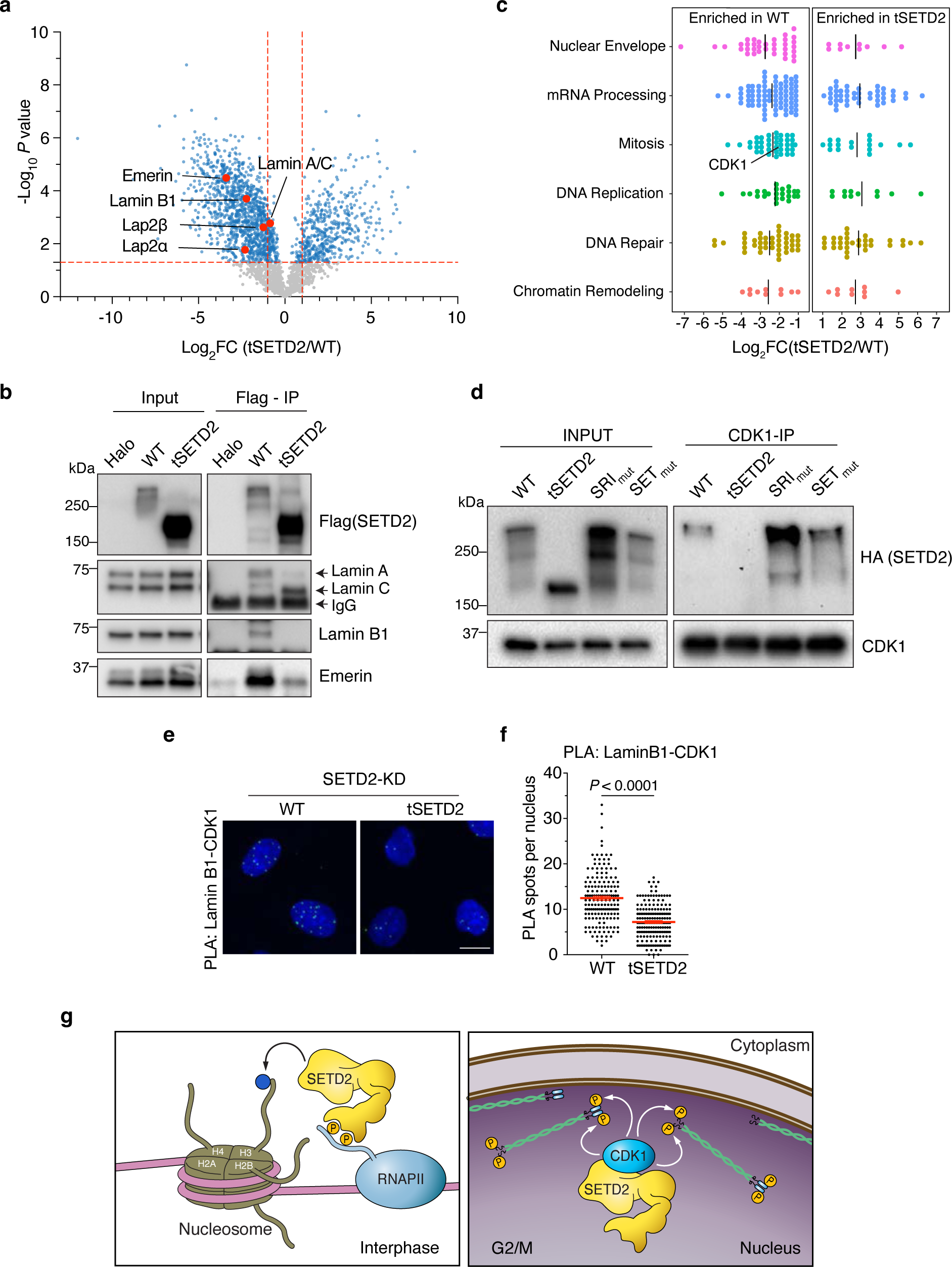
SETD2 N-terminus associates with the mitotic lamin kinase CDK1. **a**, Mass-spectrometry analysis of SETD2-APEX2 and tSETD2-APEX2 interactome. Y-axis is significance, x-axis is log2FC of tSETD2/WT normalized data. Nuclear lamina proteins highlighted in red. Blue dots are significant hits (p<0.05). **b,** Immunoblot analysis of the indicated proteins following immunoprecipitation of Flag-Halo tagged full-length WT-SETD2 or tSETD2. Flag-Halo served as control. **c,** Gene Ontology analysis of APEX2 mass-spec data from a. Relative enrichment between intact and tSETD2 mass-spec datasets of the proteins in the indicated gene ontologies. **d,** Immunoblot analysis of the indicated proteins following immunoprecipitation of endogenous CDK1 from the indicated samples. **e,** Proximity ligation assay (PLA) using anti-CDK1 and antilamin B1 antibodies in SETD2-depleted cells expressing either WT SETD2 or tSETD2. PLA pairs are in green; DNA was counterstained with DAPI. Scale bar is 10µm. **f,** Quantitative analysis of PLA pairs between CDK1 and lamin B1 shown in e. Data shown are mean ± s.e.m**. g,** Model depicting SETD2’s molecular functions in interphase (left) and G2/M (right).

The N-terminus of SETD2, encompassing the first 1322 amino acids, makes up half of the total SETD2 protein. Scattered across the first half (1322 amino acids) of SETD2 are regions of low complexity, also known as intrinsically disordered regions (IDR); the highest concentration of IDRs is located within the first 500 N-terminal amino acids (Fig. 5a and Extended data 5a). Because of their ability to promote liquid-liquid phase separation, IDRs have important functions in biology^33^. Therefore, we asked whether the IDRs and/or other specific regions within the N-terminus of SETD2 are important for its function in the maintenance of the nuclear lamina. We created a series of dox inducible N-terminal deletion mutants, namely, deletion of 1-503 a.a (region A), 504-816 a.a (region B), 817-1322 a.a (region C), and 504-1322 a.a (region B+C) (Extended data 5a). As previously observed, intact SETD2 formed distinct foci in the nuclei of interphase cells (Extended data 5b). The association of SETD2 with elongating RNAPII and splicing factors promoted us to ask whether these foci were nuclear speckles, which was confirmed by the finding that SETD2 foci co-localized with the nuclear speckle marker SRSF1^34^. Surprisingly, deletion of the entire N-terminus (tSETD2) or just region A (ΔA, 1-503 a.a) abrogated SETD2 localization to nuclear speckles. Importantly, a fusion of N-terminal region A directly to tSETD2 (i.e., ΔBC, 504-1322 a.a) restored SETD2 localization to speckles. Thus, the first 503 amino acids of SETD2 are both necessary and sufficient to localize SETD2 to nuclear speckles. In stark contrast to these observations, SETD2 lacking region A fully rescued the nuclear lamina and genome integrity defects (Extended data Fig. 5c, d), which indicated that this activity of SETD2 is embodied within the B and/or C regions of the N-terminus. Deletion of either region B or region C alone produced only minimal effects on nuclear defects, whereas deletion of both B and C regions phenocopied the loss of SETD2 (Extended data Fig. 5c, d). Thus, regions B and C are required for SETD2 to maintain nuclear lamina integrity. These results also revealed that distinct functions of SETD2 are harbored within the N-terminus (i.e., region A (1-503a.a) drives SETD2 to nuclear speckles and regions B and C (504-1322 a.a) promote maintenance of lamina integrity).

### SETD2 associates with the mitotic kinase CDK1 to coordinate lamin phosphorylation

To better understand how the N-terminus of SETD2 controls nuclear lamina integrity, we further examined the protein-protein interactions differences between WT SETD2 and tSETD2. Gene Ontology (GO) term analysis of these mass spectrometry data revealed a significant enrichment of proteins related to nuclear envelope, mRNA processing, and mitosis among WT SETD2 interactors compared to tSETD2 interactors. Surveying the list of proteins in the “mitosis” group, we noticed that cyclin dependent kinase 1 (CDK1) was significantly enriched as a SETD2 interactor compared with tSETD2 (Fig. 6c). This observation was intriguing because CDK1 is the master kinase that regulates the G2-to-M transition by phosphorylation of multiple substrates, including lamins^35^. Importantly, CDK1 phosphorylation of lamin A/C at S22, S390, S92 and lamin B1 at S23, S391, S393 during G2-M transition is a required step in disassembly of lamins and the nuclear envelope^17, 25^. The finding that this kinase also associates with the N-terminus of SETD2, a result we confirmed by several co-immunoprecipitation experiments (Fig. 6d and Extended data Fig. 5e), deepened the connection between SETD2 and lamina regulation.

The association of SETD2 with lamins and CDK1 suggested that the SETD2 N-terminus functions as a scaffold that locally co-organizes CDK1 and lamins, thereby facilitating lamin phosphorylation. If this hypothesis was correct, we would expect that deletion of the N-terminus of SETD2 would reduce the association of CDK1 with lamins during mitosis. We tested this idea by performing PLA experiments using antibodies against CDK1 and lamin B1 in SETD2-depleted cells that expressed either full-length SETD2 or tSETD2 (Fig 6 e, f). We found that cells expressing tSETD2 had a significant decrease in the interaction frequency between CDK1 and lamin B1 compared to full-length SETD2 expressing cells. Thus, the N-terminus of SETD2 is important for association of CDK1 with lamins, and this property of SETD2 may explain why the absence of SETD2 or its N-terminus results in decreased lamin phosphorylation and altered nuclear lamina dynamics – events that destabilize nuclear lamina integrity and result in genome instability (Fig. 6g).

## Discussion

Our studies reveal an unsuspected link between SETD2, the nuclear lamina, and CDK1, which is important for the integrity of the nuclear lamina and the genome. Our findings also suggest a model wherein SETD2, via its N-terminus, serves in a structural manner to facilitate the CDK1-mediated lamin phosphorylation during mitosis that contributes to nuclear lamina disassembly, a process that when improperly regulated leads to genome instability. To date, most phenotypes associated with SETD2 loss have been attributed to deficiencies in its methylation activity. However, using chemical inhibition and genetic complementation experiments, we demonstrated that SETD2 participates in a fundamental aspect of the cell cycle that is independent of its catalytic activity. Many other chromatin regulators contain large, disordered regions similar to SETD2; thus, we predict that some of these regions may be revealed to have critical non-catalytic and non-nucleosomal activities. Interestingly, a study by Schibler et al has implicated a wide number of chromatin regulators to be involved in nuclear morphology regulation; it will be exciting to determine if protein regions outside of the enzymatic activities of these chromatin regulators is involved in nuclear morphology regulation^36^.

An intriguing observation from these studies is the delineation of activities found within the N-terminus of SETD2. Although the first 500 amino acids (region A) contribute to SETD2 recruitment to nuclear speckles, its B and C regions (504-1322 a.a) act completely differently, functioning as a scaffold for CDK1 and lamins to ensure lamin phosphorylation and depolymerization. It remains to be determined how the A region of SETD2 contributes to transcription or splicing in nuclear speckles, although we hypothesize that recruitment of SETD2 to nuclear speckles, which are associated with high levels of transcription^34, 37^, may aid in the direct recruitment of RNA processing and splicing machinery to these regions. In contrast to the A region, we defined cell cycle related functions of SETD2 for the B and C regions of the N-terminus. The idea that SETD2 would have a cell cycle function in mitosis is supported by studies that show SETD2 regulates the timing of the cell cycle and that its protein level is stabilized during the G2/M phase^29^. Additionally, mammalian SETD2 is directly targeted for phosphorylation by CDK1^38^. In all, these findings paint a vital function for SETD2 that is not expected from its previously defined activities in transcription. Interestingly, the extended SETD2 N-terminus required for the nuclear lamina function we describe here is missing in less complex organisms such as yeast Set2^2, 3^. We surmise that evolutionary growth of the N-terminus of SETD2 likely arose due to a necessity for SETD2 to fulfill multiple functions as the complexity of eukaryotic cells increased.

As mentioned earlier, our findings also showed that SETD2 levels increase at G2/M, which coincides with the timing of lamin phosphorylation. This temporal regulation is likely not accidental and may point to a key G2/M function for SETD2 to tether CDK1 and lamins together to promote lamin phosphorylation. The phenotypes in the absence of SETD2 or its N-terminus are fully consistent with this idea. However, the mechanism by which SETD2 stabilization occurs in G2/M is not yet known. Michowski et al. found that the SETD2 N-terminus was directly targeted by CDK1^38^; thus, it may be that CDK1 phosphorylation of SETD2 contributes to its G2/M stability, central to the mechanism we describe herein.

Another key observation from this study is that the precise amount of SETD2 is of great importance. This fact was best illustrated with our model MEF cells that could create *Setd2* haploinsufficiency and the finding that loss of one allele resulted in dramatic defects in nuclear morphology and genome stability, even though H3K36me3 levels were unaltered (Fig. 4d, e). Because greater than 90% clear cell renal cell carcinomas (ccRCCs) have heterozygous deletions of *SETD2*, it is intriguing to speculate that the loss of SETD2 in ccRCC is a key driving event for the initiation of these tumors and perhaps other tumors that have heterozygous deletion of *SETD2*. We also note that ∼15% of ccRCCs exhibit a homozygous loss of *SETD2*, which has been suggested to be a key secondary event for this cancer population^5^. Many of these “second hit” mutations are missense mutations that occur within the SET domain. Of these, R1625C is a hotspot mutation thought to create a null allele because it renders the enzyme inactive^5^. However, we showed that this particular mutation destabilizes SETD2 protein^39^, thereby creating a hypomorph in addition to being inactive. Therefore, it is conceivable that heterozygous and homozygous *SETD2* loss are important etiologies of ccRCC by protein regulation that affects nuclear lamina stability. Finally, we note that mutations in lamins or changes in their expression lead to nuclear morphological defects, which, like mutant SETD2, can promote tumorigenesis by increased aneuploidy and genome instability^40^. Because SETD2 is frequently mutated in several cancers, especially ccRCCs, which are marked with genome stability defects, we speculate that the function of SETD2 as a tumor suppressor may be mediated largely by its control of nuclear envelop disassembly during the cell cycle.

## Acknowledgements

We thank all Strahl laboratory and Davis laboratory members for valuable input. We thank Supriya Prasant for critical comments on the manuscript, Wendy Salmon for advice and helpful suggestions related to microscopy, Kohta Ikegami for sharing *LMNA* mutant cell lines, Kimryn Rathmell for sharing *Setd2* floxed MEFs, Wesley Legant and Fariha Rahman for help with microscopy, and Zachary Mayo for help with initial experiments. We thank the UNC Proteomics Core Facility and the UNC Hooker Imaging Core Facility, which are supported in part by NCI Cancer Center Core Support Grant (2P30CA016086-45) to the UNC Lineberger Comprehensive Cancer Center. This work was supported by NIH grant to B.D.S. (GM126900).

## Author Contributions

A.K. and B.D.S. conceived the study. A.K. and J.M.M. performed all the experiments with help from L.C.C. and A.L.B. A.K. and B.D.S. analyzed data. B.B. and M.B.M. helped with initial mass-spec experiments. C.A.M. and L.E.H. performed mass spectrometry analyses. K.L. and J.A. synthesized EPZ-719. I.J.D. provided technical resources and conceptual ideas. A.K. and B.D.S. wrote the paper with input from all authors.

## Corresponding author

Correspondence requests for materials should be addressed to

## Ethics declarations

B.D.S. is a co-founder and board member of EpiCypher, Inc.

## Data Availability

Raw and unprocessed mass spectrometric data will be available at the PRoteomics IDEntifications (PRIDE) database (https://www.ebi.ac.uk/pride/).

---

**Extended Data Fig. 1.**
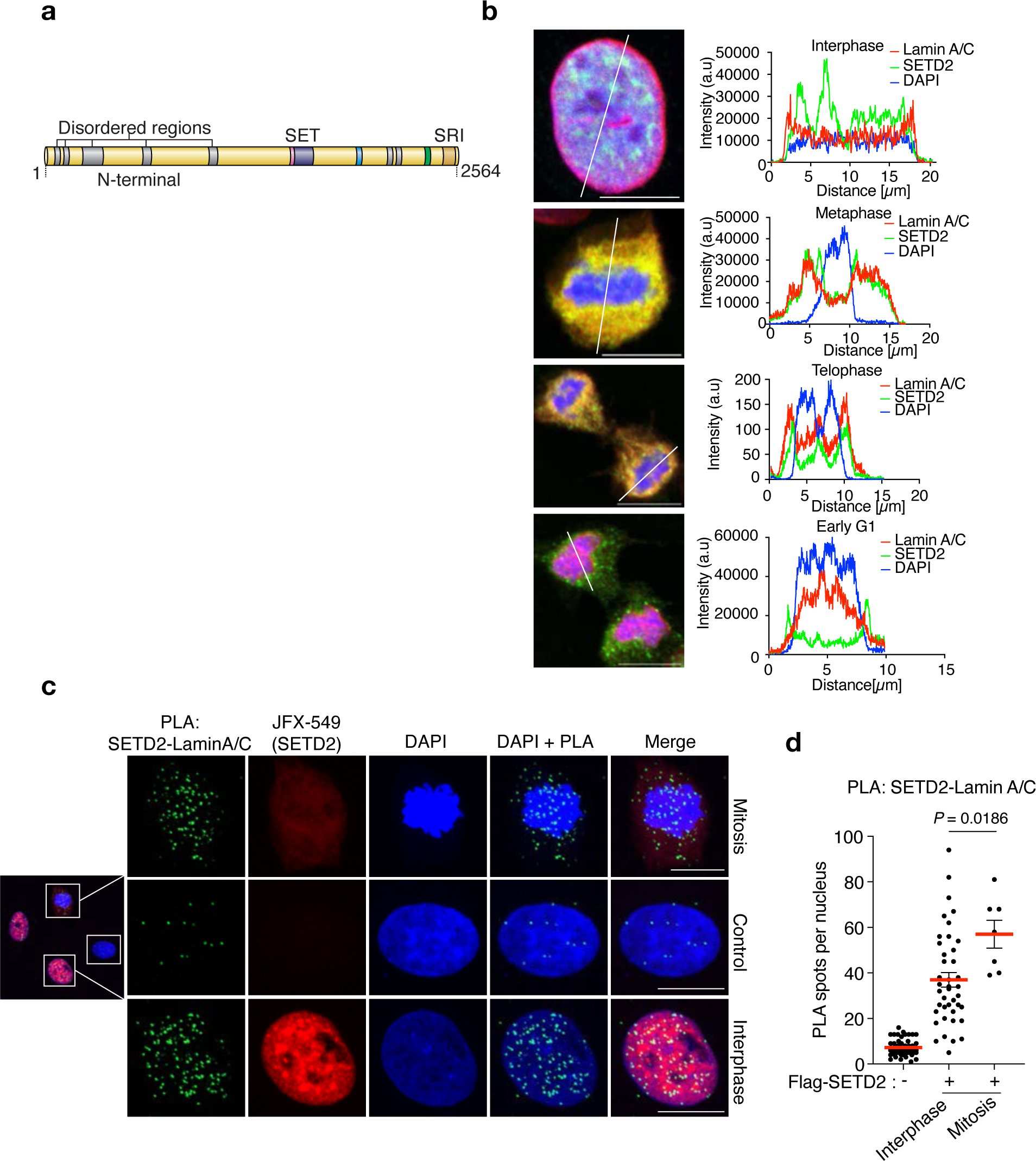

**Extended Data Fig. 2.**
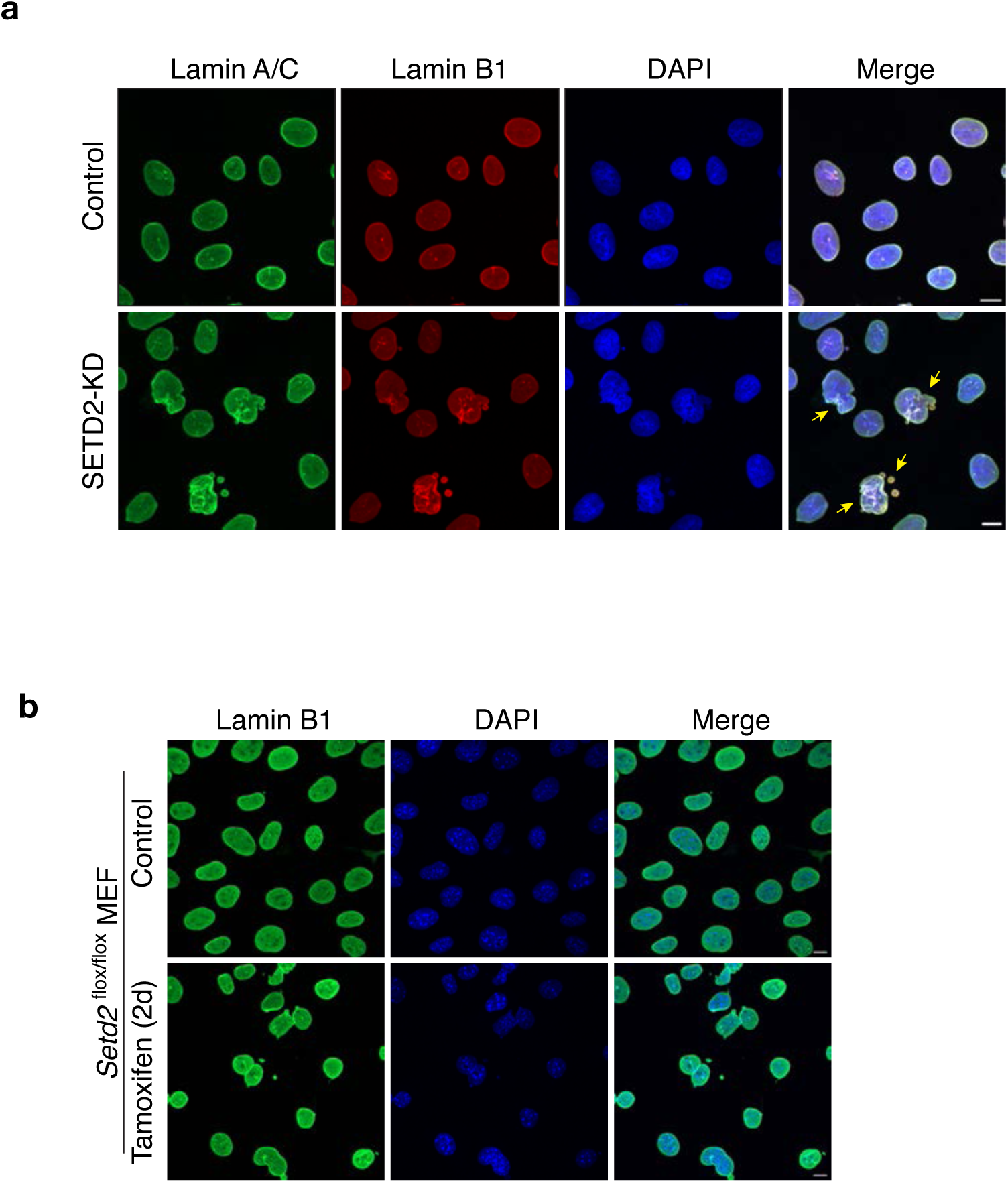

**Extended Data Fig. 3.**
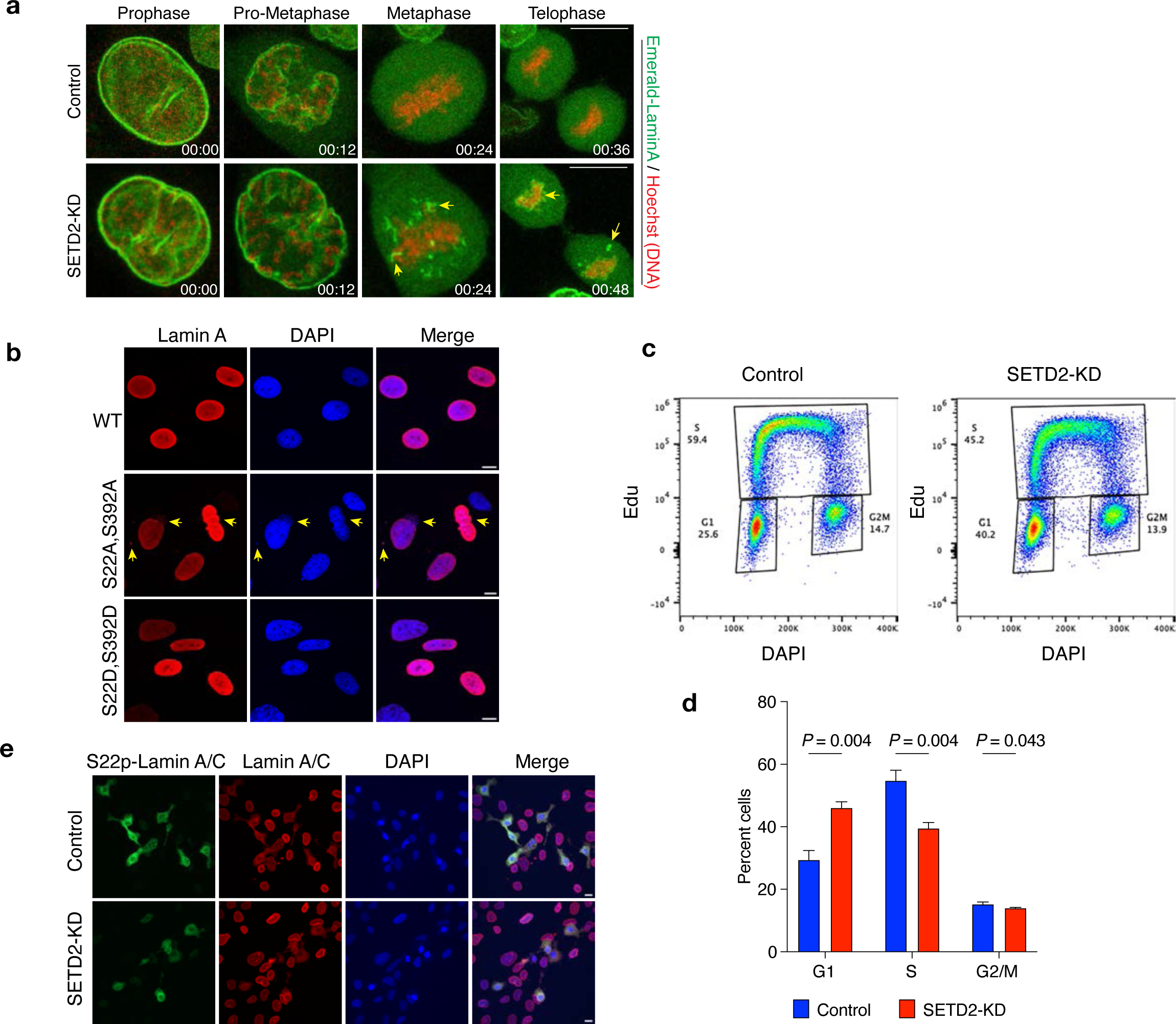

**Extended Data Fig. 4.**
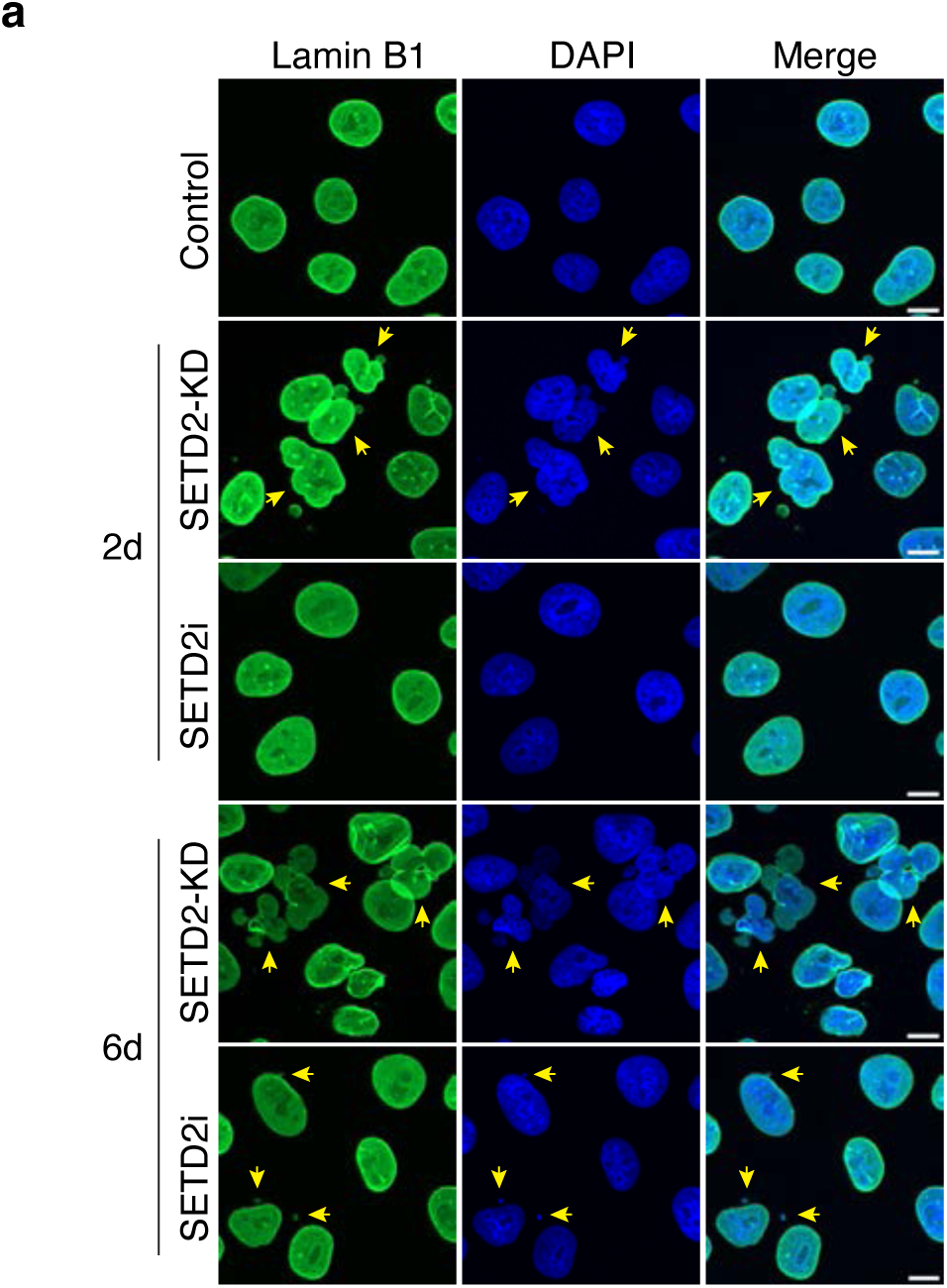

**Extended Data Fig. 5.**
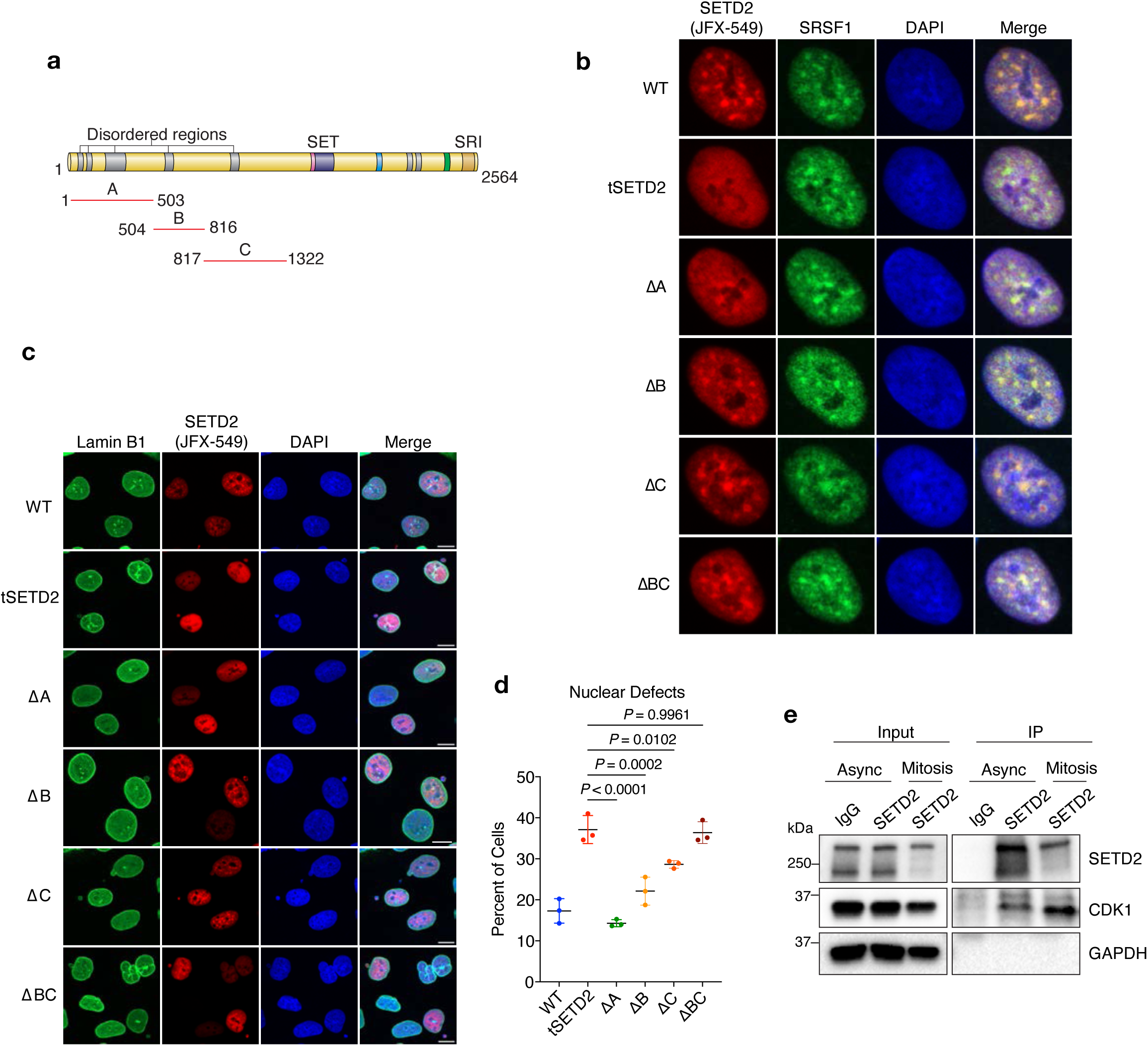

