## Supplemental_methods for "SETD2 maintains nuclear lamina stability to safeguard the genome"

### Figure Legends

**Extended Data Fig. 1 | SETD2 interacts with nuclear lamina proteins. a,** Schematic of SETD2 with its indicated domains. **b,** Fluorescence intensity profiles of SETD2, lamin A/C and DAPI in interphase or mitotic cells. Scale bar is 10 μm. **c,** Proximity ligation assay (PLA) on SETD2-KD cells expressing Halo-Flag tagged SETD2 using anti-SETD2 and anti-lamin A/C antibodies. PLA pairs are in green; SETD2 (red) visualized using fluorescent Halo ligand JFX-549. **d,** Quantitative analysis of PLAs between SETD2 and lamin A/C shown in c. DNA was counterstained with DAPI. Scale bar is 10 μm.

**Extended Data Fig. 2 | SETD2 loss leads to nuclear lamina defects. a,** Representative confocal microscopy images of control and SETD2-KD HKC cells immunostained with anti-lamin A/C and lamin B1 antibodies. DNA was counterstained with DAPI. Scale bar is 10 μm. **b,** Representative confocal microscopy images of control and *Setd2*-KO MEF cells immunostained with anti-lamin B1 antibody. DNA was counterstained with DAPI. Scale bar is 10 μm

**Extended Data Fig. 3 | SETD2 loss impairs lamin phosphorylation during G2/M. a,** Representative time-lapse images of control and SETD2-depleted HKC expressing Emerald-lamin A (green). DNA was labeled with SiR-Hoechst (red). Arrowheads point to partially depolymerized lamin filaments observed throughout mitosis. **b,** Representative images of *LMNA* KO human fibroblasts cells expressing either WT, S->A (S22A; S392A) or S->D (S22D; S392D) mutant lamin A. Cells were immunostained with lamin A/C antibody (red) and DNA was counterstained with DAPI. **c,** Representative flow cytometry analysis of control and SETD2-KD cells to determine cell-cycle distribution. Y-axis is EdU intensity and x-axis DNA content. **d,** Quantitation of flow cytometry analysis data from three independent replicates showing cell-cycle distribution of control and SETD2-KD cells. **e,** Representative images of S22phos-lamin A/C (green) and pan lamin A/C (red) from control and SETD2 depleted cells. DNA was counterstained with DAPI. All scale bars are 10 µm.

**Extended Data Fig. 4 | Nonenzymatic function of SETD2 in maintenance of nuclear integrity. a,** Representative images of control, SETD2-KD or SETD2i (EPZ-719) treated HKC cells for 2d or 6d, immunostained with lamin B1 antibody. DNA was counterstained with DAPI. Scale bar is 10 µm.

**Extended Data Fig. 5 | The N-terminus of SETD2 regulates nuclear lamina integrity. a,** Schematic of SETD2 protein with indicated regions (A, B, or C) deleted to create individual N-terminal mutants. **b,** Representative confocal images of SETD2-depleted cells expressing either WT-SETD2 or the indicated N-terminal mutants, immunostained with SRSF1 (green). SETD2 (red) visualized using Halo ligand JFX-549. **c,** Representative confocal images of SETD2-depleted cells expressing either WT SETD2 or the indicated N-terminal mutants, immunostained with lamin B1 (green). SETD2 (red) visualized using Halo ligand JFX-549. DNA was counterstained with DAPI. Scale bar is 10 µm. **d,** Quantification of total nuclear defects in HKC cells expressing either WT SETD2 or the indicated N-terminal mutants. **e,** Immunoblot analysis for the indicated proteins following immunoprecipitation of endogenous SETD2 in asynchronous or mitotic cells.

### Methods

##### Chemistry

EPZ-719 was synthesized using a modification of the published route (Scheme S1).[30]
Compound **1** was reacted with MsCl to afford **2**, which was deprotected with HCl, followed by reductive amination with *tert*-butyl (*R*)-(3-oxocyclohexyl)carbamate to give **3**. The *N*-

deprotection of **3** and amide coupling with 4-fluoro-7-methyl-1H-indole-2-carboxylic acid to afford EPZ-719. The $^{1}H$ and $^{13}C$ NMR spectra match the reported values.[30]

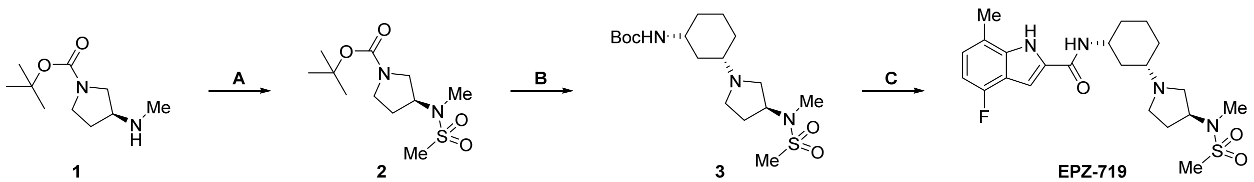


**Scheme S1. Modified synthesis of EPZ-719.** (A) MsCl, diisopropylethylamine, DCM, 91%; (B) (1) HCl/DCM/ether, then (2) *tert*-butyl (*R*)-(3-oxocyclohexyl)carbamate, $NaBH_4$, MeOH/NaOMe, 38% over 2 steps; (C) (1) HCl/DCM/ether, then (2) 4-fluoro-7-methyl-1H-indole-2-carboxylic acid, DIC, HOBt, DMAP, diisopropylethylamine, DCM, 43% over 2 steps.

**Antibodies**

SETD2 (in-house generated, 1:1000), GAPDH (Cell Signaling, 14C10, 1:5000), H3K36Me3 (Abcam, ab9050 and in-house generated, 1:5000), CDK1 (Millipore, 06-9230, 1:2000), Lamin A/C (Sigma, SAB4200236, 1:2000), Ser22 Lamin A/C Phos (Cell Signaling, 13448S, 1:1000), RNAPII RPB1 (Cell Signaling, 14958S, 1:1000), M2 FLAG (Sigma, F1804, 1:2000), Emerin (Thermo, MA5-31328, 1:2000), Lamin B1 (Abcam, ab16048, 1:2000), Lamin B1 (ProteinTech, 66095-1-Ig, 1:2000), SRSF1 (ThermoFisher, 32-4600, 1:2000). CDK1 (clone A17.1.1 Millipore-Sigma, MAB8878, 1: 1000).

**Cell culture and treatment**

Human kidney tubule epithelial cells (HKC) were obtained from Dr. Lorraine Racusen, (Johns Hopkins Hospital, Baltimore, MD) and Bj5a *LMNA* mutant fibroblast cells were a gift from Dr. Kohta Ikegami (Cincinnati Children's Hospital). HKC and Bj5a were maintained in Dulbecco's modified Eagle's medium (DMEM) supplemented with 100 U $ml^{-1}$ penicillin, 100 μg $ml^{-1}$ streptomycin and 10% fetal calf serum. *Setd2*$^{fl/fl}$ and *Setd2*$^{fl/wt}$ mouse embryonic fibroblasts (MEFs), a gift from Dr. W. Kimryn Rathmell (Vanderbilt University), were maintained in phenolred free DMEM supplemented with 100 U $ml^{-1}$ penicillin, 100 μg $ml^{-1}$ streptomycin and 10% fetal calf serum. All cultures were maintained at 37°C and 5% $CO_2$. To induce *Setd2* deletion, MEFs were treated with 3 μM 4-hydroxytamoxifen (4-OHT) for 48 h. HKC cells were synchronized using double-thymidine block method by treating cells with 2 mM thymidine for 16 h, followed by release for 8 h and re-treatment with 2 mM thymidine for 16 h to bring cells to G1/S boundary. Cells were then released for appropriate times based on the downstream analyses. For mitotic arrest, cells were treated with 50 ng/ml of nocodazole for 12 h.

**Cloning/Plasmid generation**

shRNA against SETD2 (TRCN0000237839) was cloned into Tet-pLKO-puro vector (Addgene, 21915). Dox-inducible full-length SETD2 constructs were generated by Gibson assembly of the N-terminus (1-1322 a.a), the C-terminus (1323-2564 a.a) and Halo-3xFLag (gBlock, IDT) into Xlone-GFP (Addgene, 96930) following KpnI/SpeI digestion to excise EGFP cDNA. Generation of the C-terminus (also referred to as tSETD2) and its mutant derivatives (R1625C, R2510H) were previously described[39]. Mutants consisting of N-terminal truncations or deletions of regions A, B and/or C (Extended Data Fig. 5a) were also generated by Gibson assembly.

**Generation of stable cell lines**

HKC shSETD2-TetOn cells were generated by infecting cells with lentivirus containing the dox-inducible shSETD2 construct, followed by selection with puromycin (1 μg/ml). To induce SETD2 knockdown (KD), HKC-shSETD2-TetOn cells were treated with 500 ng/ml of doxycycline for 48-72 h. To generate SETD2 complementation cell lines, HKC-shSETD2-TetOn cells were nucleofected with a transposase expression vector (System Biosciences, PB210PA) and Xlone-piggyBac plasmid containing dox-inducible SETD2 transgene (WT or mutant), followed by selection with increasing concentration of blasticidin (2-10 μg/ml) over a period of two weeks.

Cells were treated with doxycycline at 500 ng/ml for 5-6 days to allow for simultaneous knockdown (KD) of endogenous SETD2 and expression of the exogenous SETD2 transgene before downstream analyses.

**Co-immunoprecipitation**

Cells were resuspended in ice-cold lysis buffer containing 50 mM Tris-HCl pH 7.5, 150 mM NaCl, 10% glycerol, 0.25% Triton-X100, 1 mM $MgCl_2$, 1 mM PMSF, 1 U/μl Universal Nuclease (Pierce) supplemented with protease and phosphatase inhibitors. Lysates were incubated on rotation for 2 hours followed by brief sonication and centrifugation at 13,000g for 20 min. Lysates were pre-cleared using equilibrated Protein-G Sepharose beads for 1 h. Pre-cleared lysates were incubated with 2-4 μg of indicated antibody conjugated Protein-G Dynabeads (Life Technologies) for 4 hours or overnight. Beads were gently centrifuged, separated on magnet and washed five times with lysis buffer (with 1mM EDTA) for 10 min each wash. Immunoprecipitated proteins were eluted by boiling the beads in SDS sample loading buffer for 5 min at 95$^o$C.

**Immunofluorescence**

Cells grown on glass coverslips were fixed in 4% formaldehyde (methanol-free) in PBS for 15 min at room temperature. Cells were washed three times with PBS for 5 min each, followed by permeabilization with 0.5% Triton in PBS for 10 min. Cells were blocked in 2% BSA, 1% normal goat serum (NGS) in PBS for 30 min at RT prior to antibody incubation for 1 hour at RT. Cells were washed thrice with PBS for 5 minutes each, followed by secondary antibody incubation (Alexa Fluor, Life Technologies) at 1:1000 for 1 hour at RT. Cells were washed with PBS followed by DAPI staining and mounting on glass slides with ProLong Gold Antifade mounting media (Life Technologies). For visualization of Halo tagged SETD2 constructs, cells were treated with 100nM JFX-549 (Janelia Farms) ligand for 20 min before fixing.

**Confocal microscopy**

Images were acquired using Zeiss LSM 800 confocal microscope using either a 40x/1.3NA oil or 63x/1.4NA oil objective. Z-stacks of 0.5μm slices were collected and maximum intensity projections were used for analysis and display. For colocalization analyses, fluorescence intensity profiles were extracted for all fluorophores from the same Z-axis plane using the Zen Blue software and graphed using Prism 9. Images were processed and analyzed using ImageJ (version 2.3.0) and CellProfiler (version 4.2.4).

**Live cell microscopy**

HKC-shSETD2-TetON cells expressing Emerald-lamin A were plated in an 8-well chambered cover glass (Cellvis). Cells were treated with dox one day prior to synchronization using double thymidine block and released for 4 hours in the presence of 100nM SiR-Hoechst (Spirochrome) prior to imaging. Images were acquired using Zeiss LSM 880 confocal microscope with a 40x/1.3NA oil immersion objective. Z-stacks of 1μm slices were collected every 4 or 10 min intervals for 6-8 hours. Images were processed and analyzed using ImageJ (version 2.3.0).

**Proximity Ligation Assay (PLA)**

PLA was performed according to the manufacturer's protocol (Duolink PLA kit, Sigma) with minor modifications. Briefly, after fixation and permeabilization, cells were treated with previously described blocking solution[41] (10% NGS, 2% BSA, 5% sucrose in PBS) for 45 min, followed by primary antibody incubation in blocking solution for 1 hour at RT. Cells were washed and treated with plus and minus probes diluted in blocking solution for 1 hour at $37^{o}$C. Probes were ligated for 30 min at $37^{o}$C followed by *in situ* PCR amplification using fluorescent oligonucleotides (green or red) for 120 min at $37^{o}$C. Cells were counterstained with DAPI and mounted using ProLong Gold Antifade mounting media (Life Technologies). Images were acquired on a Zeiss 880 confocal microscope with 63x/1.4 oil immersion objective. Images were processed using ImageJ and PLA pairs were quantified using CellProfiler (version 4.2.4)[42].

**Image analysis**

For PLA analyses, a published pipeline for foci counting from cellprofiler.org was adapted to quantify PLA spots. Briefly, nuclei were segmented based on DAPI signal. Next, PLA foci were detected with 4-30 pixels limit and global thresholding. PLA objects were related to parent nuclei objects and per object counts were calculated and exported as spreadsheet. To detect SETD2 expressing cells, an additional step was added to detect SETD2 intensity as secondary object per nucleus. For live-cell image analysis in Fig. 3c, cells were manually tracked and fluorescent intensities were extracted using ImageJ. Briefly, ROIs were drawn encompassing Emerald-lamin A signal for every frame from the last metaphase frame immediately preceding anaphase onset to the end of mitosis (typically 10 frames of 4 min intervals) and mean fluorescence intensity (MFI) was extracted for Emerald-lamin A. MFI of every frame was normalized to the first frame (last metaphase frame, set as 0 min) and the time taken to reach maximum MFI was determined for each cell analyzed. Maximum MFI of Emerald-lamin A was indicative of complete lamina reassembly.

**Cell cycle analysis**

Cells were treated with 10 μM EdU for 20 min before collection. Cells were trypsinized, washed with PBS once and pelleted. Cells were fixed in 4% formaldehyde in PBS for 15 min. After permeabilization and blocking in 1% BSA, click reaction was performed to detect EdU incorporation. Briefly, cells were incubated in PBS with 1mM $CuSO_4$, 100 mM ascorbic acid and 1 μM Alexa-647-azide for 30 min at RT in the dark. After washes, cells were resuspended in PBS-BSA containing DAPI and 100 μg/ml RNase A overnight. Flow cytometry was performed on Attune

Nxt flow cytometer (Thermo Fisher) and data were processed and analyzed using FlowJo (BD Biosciences).

**Western blot**

Cells were lysed in SDS sample buffer. Lysates were sonicated and boiled for 10 min at 95$^{o}$C. Proteins were separated on a 4-15% gradient SDS-PAGE gel (BioRad) and proteins were subsequently transferred onto a PVDF membrane. Membranes were blocked in 5% non-fat milk in TBST and incubated with antibodies overnight in milk. Membranes were washed and incubated with secondary antibodies for 1 h prior to ECL treatment and signal detection using ChemiDoc imaging system (BioRad).

**Proximity biotinylation and affinity purification**

HKC cells stably integrated with dox-inducible WT-SETD2-APEX2 or tSETD2-APEX2 were induced with dox for three days prior to proximity labeling and affinity purification. Corresponding cells without dox treatment were used at controls. Proximity labeling and streptavidin AP was performed as previously described[14] with a few modifications. Briefly, cells were pulsed with 500 μM biotin-phenol for 30 min at 37$^{o}$C. Biotinylation was performed with 1 mM $H_2O_2$ for 1 min and reaction was stopped by rinsing cells with quencher solution (10 mM sodium ascorbate, 5 mM Trolox, and 10 mM sodium azide in 1xPBS). Cells were resuspended in lysis buffer (50 mM Tris-HCl pH 7.5, 200mM NaCl, 10% glycerol, 0.25% Triton-X100, 1 mM $MgCl_2$, 1 mM PMSF, 1 U/μl Universal Nuclease (Pierce) supplemented with protease and phosphatase inhibitors). Lysates were incubated on rotation for 2 hours followed by brief sonication and centrifugation at 13,000g for 20 min. Protein lysates were then dialyzed against lysis buffer overnight with two changes of buffer to remove excess free biotin-phenol. Lysates were centrifuged at 13,00g for 20 min and incubated with equilibrated streptavidin magnetic beads (Pierce) for 4 h. Beads were washed and proteins were eluted off the beads as described previously[14]. Eluted proteins were separated on

4-15% gradient SDS-PAGE gel for 1 cm or until the 25kDa band was visible. Gel was stained with colloidal blue staining (Thermo, LC6025) and lanes were excised and the proteins were reduced with 5 mM DTT, alkylated with 15 mM iodoacetamide, and in-gel digested with sequencing grade trypsin (Promega) overnight at 37˚C. Peptides were extracted, desalted with C18 Desalting Spin Columns (Thermo) and dried via vacuum centrifugation. Peptide samples were stored at -80˚C until further analysis.

**Mass-spectrometry analysis**

*SETD2 interactome analysis*: Streptavidin pulldown samples (n=3) were analyzed by LC/MS/MS using an Easy nLC 1200 coupled to a QExactive HF mass spectrometer (Thermo Scientific). Samples were injected onto an Easy Spray PepMap C18 column (75 μm id × 25 cm, 2 μm particle size) (Thermo Scientific) and separated over a 90-minute method. The gradient for separation consisted of 5–45% mobile phase B at a 250 nl/min flow rate, where mobile phase A was 0.1% formic acid in water and mobile phase B consisted of 0.1% formic acid in 80% ACN. The QExactive HF was operated in data-dependent mode (DDA) where the 15 most intense precursors were selected for subsequent fragmentation. Resolution for the precursor scan (m/z 350–1700) was set to 60,000, while MS/MS scans resolution was set to 15,000. The normalized collision energy was set to 27% for HCD. Peptide match was set to preferred, and precursors with unknown charge or a charge state of 1 and ≥ 7 were excluded.

*Lamin A/C and Lamin B1 phosphorylation analysis***:** Lamin A/C and lamin B1 were immunoprecipitated from control and shSETD2 HKC cells as described above. Immunoprecipitated proteins were eluted by boiling and separated on 4-15% gradient gel. Gel bands corresponding to Lamin A/C or Lamin B1 were excised and reduced, alkylated, and digested as described above. Peptide samples (n=3) were analyzed by LC/MS/MS using an Ultimate 3000 coupled to an Exploris 480 mass spectrometer (Thermo Scientific). Samples were injected onto an IonOpticks Aurora series 2 C18 column (75 µm id × 15 cm, 1.6 µm particle size) and separated over a 65-minute method. The gradient for separation consisted of 2-40% mobile phase B at a 250 nl/min flow rate, where mobile phase A was 0.1% formic acid in water and mobile phase B consisted of 0.1% formic acid in 80% ACN. The Exploris 480 was operated in data-dependent mode (DDA) where duty cycle was set to 1.5 sec. Resolution for the precursor scan (m/z 375–1500) was set to 120,000, while MS/MS scans resolution was set to 15,000. The normalized collision energy was set to 30% for higher collision dissociate (HCD). Monoisotopic peak determination was set to 'peptide' and precursors with unknown charge or a charge state of 1 and >5 were excluded. For targeted analysis, Lamin A/C samples were analyzed on an Easy nLC 1200 coupled to a QExactive HF mass spectrometer (Thermo Scientific). Samples were injected onto an Easy Spray PepMap C18 column (75 µm id × 25 cm, 2 µm particle size) (Thermo Scientific) and separated over a 60-minute method. The gradient for separation consisted of 5-40% mobile phase B at a 250 nl/min flow rate, where mobile phase A was 0.1% formic acid in water and mobile phase B consisted of 0.1% formic acid in 80% ACN. Targeted parallel reaction monitoring (PRM) method was developed for a specific Lamin A/C peptide spanning S22 (unphosphorylated and phosphorylated versions). The QExactive HF was operated in MS1 and PRM mode. Resolution for the precursor scan (m/z 400–1200) was set to 60,000 with an AGC target set to $1e^6$ and a maximum injection time set to 200 ms. PRM scans (15,000 resolution) consisted of HCD set to 27; AGC target set to $1e^6$; maximum injection time set to 100 ms; isolation window of 2 Da.

**MS data analysis**

*SETD2 interaction analysis:* Raw data files were processed using MaxQuant version 1.6.15.0 and searched against the reviewed human database (containing 20,350 entries), appended with a contaminants database, using Andromeda within MaxQuant. Enzyme specificity was set to trypsin, up to two missed cleavage sites were allowed, and methionine oxidation and N-terminus acetylation were set as variable modifications, with carbomidomethylated cysteines set as a static modification. A 1% FDR was used to filter all data. Match between runs was enabled (5 min match time window, 20 min alignment window), and a minimum of two unique peptides was required for label-free quantitation using the LFQ intensities. Perseus was used for further processing[43]. Only proteins with >1 unique+razor peptide were used for LFQ analysis. Proteins with 50% missing values were removed and missing values were imputed from normal distribution within Perseus. Log2 fold change (FC) ratios were calculated using the averaged Log2 LFQ intensities of $\frac{WT_{DoxON}}{WT_{DoxOFF}}$ (for WT-SETD2-APEX2, Fig.1b). For Fig. 6a, difference between WT-SETD2-APEX2 and tSETD2-APEX2 was calculated as such $log2\,(\frac{tSETD2_{DoxON}}{tSETD2_{DOxOFF}}) - log2\left(\frac{WT_{DoxON}}{WT_{DoxOFF}}\right)$. Students t-test performed for each pairwise comparison, with p-values calculated. Proteins with significant p-values (<0.05) and Log2 FC >1 were considered biological interactors.

Gene Ontology analyses were conducted using DAVID[44]. Genes were searched against UP_KW_BIOLOGICAL_PROCESS, UP_KW_CELLULAR_COMPONENT, UP_KW_MOLECULAR_FUNCTION, GOTERM_BP_DIRECT, GOTERM_CC_DIRECT and GOTERM_MF_DIRECT. GO terms used are described in Table 1. Figures 1c, Fig. 6c were created using tidyverse[45] and ggplot2 packages in R (version 4.0.4).

*Lamin phosphorylation analysis:* DDA and PRM raw data files were processed using Proteome Discoverer version 2.5 (Thermo Scientific). Peak lists were searched against a reviewed Uniprot human database, appended with a common contaminants database, using Sequest. The following parameters were used to identify tryptic peptides for protein identification: 10 ppm precursor ion mass tolerance; 0.02 Da product ion mass tolerance; up to two missed trypsin cleavage sites; (C) carbamidomethylation was set as a fixed modification; (M) oxidation, (S,T,Y) phosphorylation were set as variable modifications. Peptide false discovery rates (FDR) were calculated by the Percolator node using a decoy database search and data were filtered using a 1% FDR cutoff. The PhosphoRS node was used to localize phosphorylation sites within the peptides. Peak areas were extracted using the Minora node, and log2 fold change (FC) ratios of each peptide were calculated (shSETD2/Control).

**Statistical analysis and reproducibility**

All experiments were conducted in $n = 3$ independent biological replicates except for Fig 3i, 4b, 4e, 5h, 5i where $n = 2$ independent biological replicates were performed as indicated in the figure legends. Statistical tests were chosen based on normality of the data. Data were tested for normality prior to statistical analyses. For normally distributed data, unpaired, two-tailed *t*-tests were performed with mean ± s.d indicated in the figure. For data without Gaussian distribution such as in Fig 2c, 2d, two-tailed non-parametric Mann-Whitney *U* test was performed with median indicated in the figure. Similarly, for comparison of multiple groups with non-normal data, Kruskal-Wallis with Dunn's multiple comparisons test was performed as indicated in the figure legend of Fig 5f, 5g. For comparison of multiple groups with normal data in Fig 5d, 5e, one-way ANOVA with Dunnett's multiple comparisons test was performed. No statistical methods were used to predetermine sample size. Cell numbers are provided in the figure legends and are based on what is generally used in the field. All non-parametric tests were performed on independent biological samples in aggregate. All statistical analyses were performed on GraphPad PRISM 9.

Table 2: GO terms from Fig. 1c

| GO Term | Figure Label |
|---|---|
| KW-0498~Mitosis | Mitosis |
| GO:0005635~nuclear envelope | Nuclear Envelope |
| GO:0006281~DNA repair | DNA repair |

| | |
|---|---|
| KW-0507~mRNA processing | mRNA processing |
| GO:0006260~DNA replication | DNA replication |
| GO:0000785~chromatin | Chromatin |
| KW-0804~Transcription | Transcription |
